# DnaK response to expression of protein mutants is dependent on translation rate and stability

**DOI:** 10.1101/2021.09.29.462496

**Authors:** Signe Christensen, Sebastian Rämisch, Ingemar André

## Abstract

Chaperones play a central part in the quality control system in cells by clearing misfolded and aggregated proteins. The chaperone DnaK acts as a sensor for molecular stress by recognising short hydrophobic stretches of misfolded proteins. As the level of unfolded protein is a function of protein stability, we hypothesised that the level of DnaK response upon overexpression of recombinant proteins would be correlated to stability. Using a set of mutants of the *λ*-repressor with varying thermal stabilities and a fluorescent reporter system, the effect of stability on DnaK response and protein abundance was investigated. Our results demonstrate that the initial DnaK response is largely dependent on protein synthesis rate but as the recombinantly expressed protein accumulates and homeostasis is approached the response correlates strongly with stability. Furthermore, we observe a large degree of cell-cell variation in protein abundance and DnaK response in more stable proteins.

## Introduction

The production of functional proteins typically requires folding into well-defined three-dimensional structures. However, protein synthesis can also lead to accumulation of unfolded, misfolded, or aggregated protein due to their marginal stability. As aggregated and misfolded proteins can have toxic effects on the cell there is a substantial fitness cost associated with them. To reduce the harm caused by aberrant proteins all living organisms have developed a sophisticated protein quality control (PQC) network. The PQC is a collection of chaperones and proteases that assists with refolding of misfolded proteins, protects from irreversible aggregation by formation of inclusion bodies or, eventually, direct misfolded or aggregated proteins for degradation. Due to the ubiquitous nature of the PQC the involved proteins are required to be highly promiscuous. E.g., hsp70 (known as DnaK in *E. coli*) binds to hydrophobic stretches approximately five residues long, that are exposed upon misfolding [1]. These short hsp70/DnaK-binding motifs are present in most proteins and allow for recognition of unfolded or misfolded proteins, in a matter largely independent of sequence and structure. Recognition of unfolded and misfolded proteins by DnaK is one of the major regulators of the transcription factor *σ*^32^ and hence PQC network in *E. coli*, although *σ*^32^ is also regulated by e.g. GroEL/S [2]. Under low stress conditions DnaK binds to *σ*^32^ targeting it for degradation by the metallo-protease FtsH [3]. In response to stress, DnaK binds preferentially to misfolded protein instead of *σ*^32^ allowing *σ*^32^ to act as a transcription factor for expression of DnaK and other PQC proteins. Other proteins in the PQC network that are regulated by *σ*^32^ includes chaperones involved in protein folding and disaggregation (ClpB, DnaJ, GroEL and GroES), cytosolic proteases that degrade proteins (ClpP, ClpX and Lon), as well as the membrane-bound FtsH involved in the regulation of *σ*^32^ levels.

Whereas it is well-established that misfolded proteins trigger a response from the PQC the molecular properties of the client proteins that impact its regulation is not fully understood. One open question is whether this response is primarily determined by the concentration of unfolded protein, or if there is a sequence dependent component to the response beyond chaperone recognition sequences. The lower the stability of a protein translated within the cell, the less will be found in the folded native state. Consequently, if the PQC is primarily dependent on concentration of unfolded or misfolded protein a correlation between protein stability and PQC response should be expected. Whether such simple relationship exists is not clear, although upregulation of GroEL, c62.5 (hsp90 homolog) and DnaK [4, 5], has been observed upon introduction of mutations that destabilize wildtype sequences. A combination of computational models and *in vitro* studies performed by Sekhar et al. suggest that DnaK binds preferentially to unstable proteins at folding equilibrium, but to slow-folding proteins while folding[6]. As DnaK can interact both co-translationally with the unfolded nascent peptide chain and with misfolded protein [7] a similar process could occur *in vivo*. However, no *in vivo* studies of the correlation between folding rates and DnaK response has been presented.

Another central question that is not fully understood is how the chaperone response is regulated by protein production. Overexpression of proteins in *E. coli* results in increased levels of chaperones [8, 9], but it is not clear to what extent this response is controlled by the level of protein translation or simply the abundance of misfolded/unfolded proteins in the cytosol. There is a negative correlation between the level of expression of a protein in *E. coli* and its dependency on DnaK for folding [10]. Thus, proteins that are difficult to fold might express at lower levels. This indicates that the cell has limited capacity to process misfolded/unfolded proteins and perhaps also a limited range to regulate the chaperone response. Mutations does not only affect the folding stability and kinetics of proteins, but also the level of expression. Therefore, to understand the regulation of the chaperone response also the protein expression and abundance must be considered.

The presence of misfolded/unfolded proteins carries a fitness cost to the cell [11-14]. By modifying the abundance of misfolded proteins, chaperones can modify the rate of protein sequence evolution [10, 15-21]. Hence, chaperones, like DnaK, can enable proteins to evolve faster. However, there is still an associated fitness cost as triggering the PQC response consumes resources; both directly by consumption of ATP and by occupying the synthesis machinery for production of chaperones and proteases forming the PQC network. In theory, the interaction of proteins with the PQC network can also impact which sequences are fixated during evolution. Protein variants triggering a large PQC response would be expected to have lower fitness and thus selected against. In light of this, information about how molecular variation impacts DnaK interaction and the PQC can provide valuable insight into the role of the PQC on protein evolution.

As summarised above the components of the PQC system and the regulation network are well described, but it has yet to be studied in detail how the biophysical properties of the client protein impact the chaperone response. Thus, in this study we investigated the effect of molecular variation on the PQC. More specifically, we focused on the following questions: (i) What is the relationship between protein stability and DnaK response? (ii) Is the DnaK response controlled by the level of protein translation, the amount of unfolded protein or both? (iii) How much does the PQC response vary with small variation in biophysical properties of protein variants? (iv) How much does the PQC response vary between individual cells? We used superfolder Green Fluorescent Proteins (sfGFP) expressed from the stress-inducible DnaK promoter to evaluate the effect on the PQC response upon introducing stability-altering point mutations to the *λ* repressor construct, N102LT. We found that induction of the DnaK promoter is dependent on protein translation rates early after onset of recombinant protein synthesis, but dependent on protein stability when homeostasis is reached. We further found that protein stability has a significant impact on accumulation of protein in the cell due to increased degradation of proteins with low stability and that the response varies substantially between cells. The investigation also indicates how the fluorescence system can be used as a high-throughput stability assay.

## Results

### Monitoring protein accumulation and DnaK response with fluorescent reporters

To study how the biophysical properties of proteins impact the stress response in *E. coli*, we constructed a reporter plasmid system to simultaneously monitor the abundance of recombinantly expressed protein and chaperone response upon recombinant expression (Figure 1A). It contains two fluorescent reporters: (i) The red fluorescent protein tagRFP is fused to the N-terminal of a protein of interest (POI), and expression of the POI-tagRFP fusion protein is controlled by the IPTG inducible *lac* promoter. The fusion protein allows for monitoring the abundance of recombinant POI in the cell. In addition, the folding state of the POI could impact the folding of tagRFP and therefor acts as an additional stability reporter, as previously shown for GFP-tagged proteins [22-24]. (ii) Expression of Superfolder Green Fluorescent Protein (sfGFP) is controlled by the DnaK promoter to enable monitoring of the stress response in individual cells. The PQC network consists of many different components, thus the GFP signal gives only a partial picture of the overall state. But since DnaK is a central component in the response to folding stress we consider it a meaningful proxy for the PQC network. However, it is worth noticing that synthesis and degradation of PQC proteins are tightly regulated in the cell [25-27]. As the degradation of GFP is decoupled from the degradation of DnaK the GFP signal does not report on the current level of DnaK in the cell, but rather the accumulated amount of DnaK produced until that point. However, by evaluating the rate of GFP production – which is correlated to the rate of DnaK synthesis – we can measure when the DnaK response is triggered. The protein abundance reporter (tagRFP) and the DnaK reporter (sfGFP) are located on the same plasmid. This enables normalization of the chaperone signal with protein abundance so that cell-cell variation and variation in plasmid copy numbers can be accounted for. It is important to note that even though a DnaK promoter is utilised for expression of GFP, the endogenous DnaK promoter is still in place ensuring a functional PQC network in the cells.

**Figure 1.**
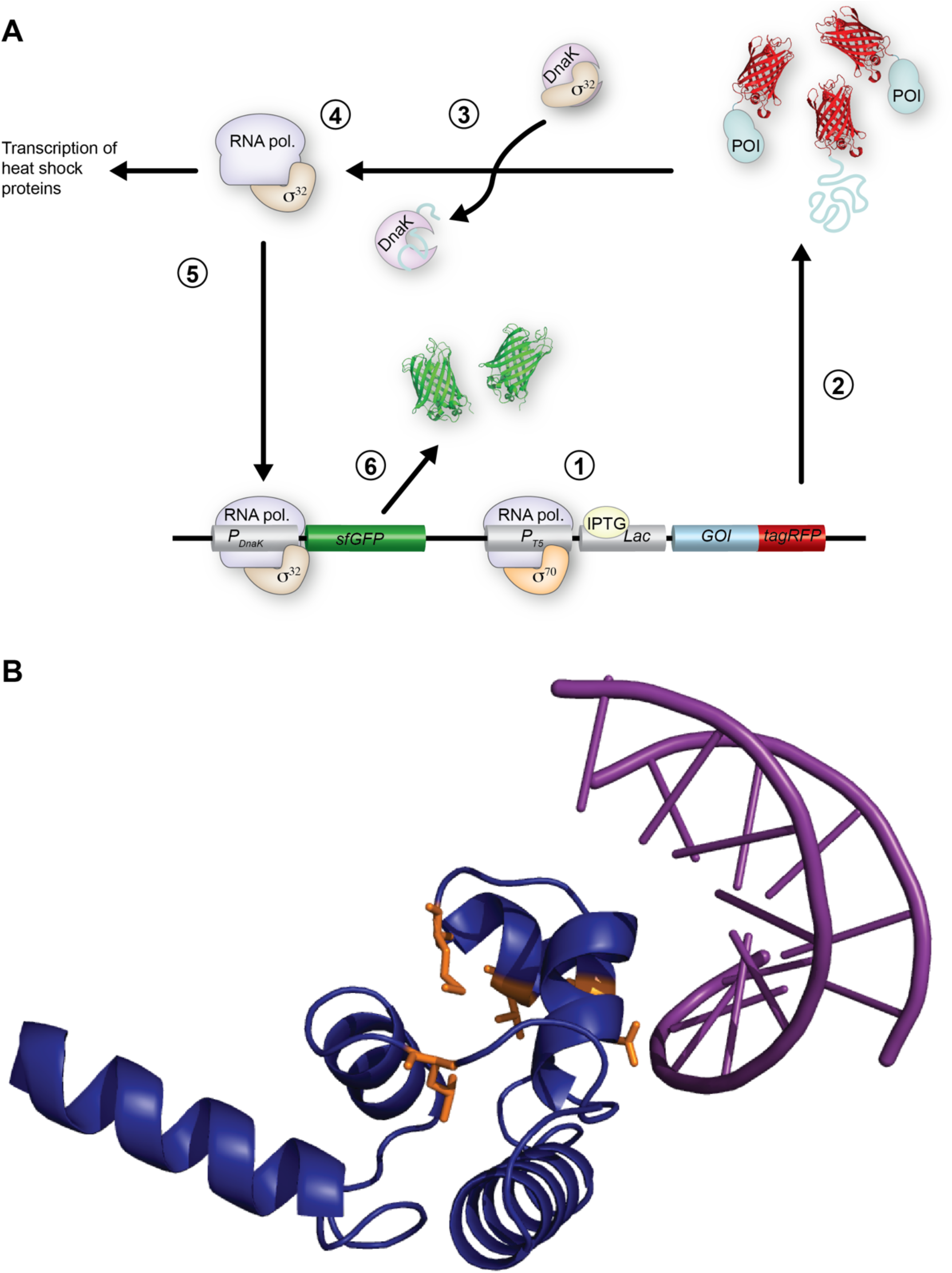
A fluorescent-based reporter system for protein abundance and DnaK response. **A)** Overview of reporter system: 1. Addition of IPTG allows for transcription of the GOI(Gene of Interest)-tagRFP by RNA polymerase in complex with the transcription factor *σ*^70^. 2. Following transcription of GOI-tagRFP the protein of interest, POI-tagRFP is synthesised. 3. DnaK releases the transcription factor *σ*^32^ and binds to unfolded/misfolded POI-tagRFP. 4. *σ*^32^ binds to RNA polymerase. 5. Binding of *σ*^32^ to RNA polymerase allows for transcription of endogenous PQC proteins, such as DnaK. The RNA polymerase - *σ*^32^ complex will also bind to the DnaK promoter on the reporter plasmid and transcribe the *sfGFP* gene. 6. Following transcription of the *sfGFP* gene, the protein sfGFP will be synthesised. **B)** The N-terminal domain of the *λ* repressor (blue) bound to DNA (purple). Mutated residues shown as sticks (orange). PDB:1lmb [28].

As a model system for investigating the correlation between DnaK response and protein stability we use several variants of the *λ* repressor construct, N102LT, with different thermodynamic stability. N102LT is a 128 amino acid long protein consisting of the N-terminal domain of the *λ* repressor extended by the C-terminal domain of the protein arc that protects the fusion protein against proteolysis [29]. It has previously been used to study the heat shock response in *E. coli* [5, 30-33]. To separate effects caused by changes in stability from effects caused by site-specific structural factors it is beneficial to also evaluate variants with mutations at the same position. In this study we therefor included seven variants with single point mutations in four different positions: Q33, V36, M40 and L57 with *in vitro* melting temperatures from below 10 °C to 61 °C in addition to wild-type N102LT (Tm = 55°C). Position L57, M40 and V36 are part of the hydrophobic core, whereas Q33 is surface exposed and is part of the cognate DNA recognition (Figure 1B). Mutations in position L57 and M40 resulted in four variants (L57C, L57G, L57P and M40A) with reduced thermostability compared to the wt sequence, while V36I results in increased stability. Mutating the surface accessible Q33 to either serine (Q33S) or tyrosine (Q33Y) retains or increases thermostability, respectively.

Changes in the ability of the N102LT variants to bind DnaK could potentially impact the observed stress response. To check whether the introduced mutations cause significant changes to the DnaK binding properties we use the ChaperISM algorithm to predict DnaK binding properties for the eight variants [34] (S1). N102LT wt is predicted to have three DnaK binding sites. Introduction of the Q33Y mutation increases the DnaK binding propensity slightly around position 33, but without introducing a new DnaK binding site. Mutating position L57 reduces the DnaK binding propensity, but without disrupting a DnaK binding site.

### Point mutations in N102LT results in differences in accumulation of protein, DnaK response and cell morphology

To monitor changes in protein abundance and DnaK response the N102LT variants were expressed from the reporter plasmid in *E. coli* and grown in liquid media. Synthesis of the N102LT variants was induced with IPTG during exponential growth. Flow cytometry was employed to simultaneous monitor the abundance of protein (red fluorescence), DnaK transcription (green fluorescence), as well as morphological changes (forward and side scatter) on single-cell level. In addition to analysis by flow cytometry, cell growth was monitored by optic density at 600 nm (OD_600_) (Figure 2A, S2).

**Figure 2.**
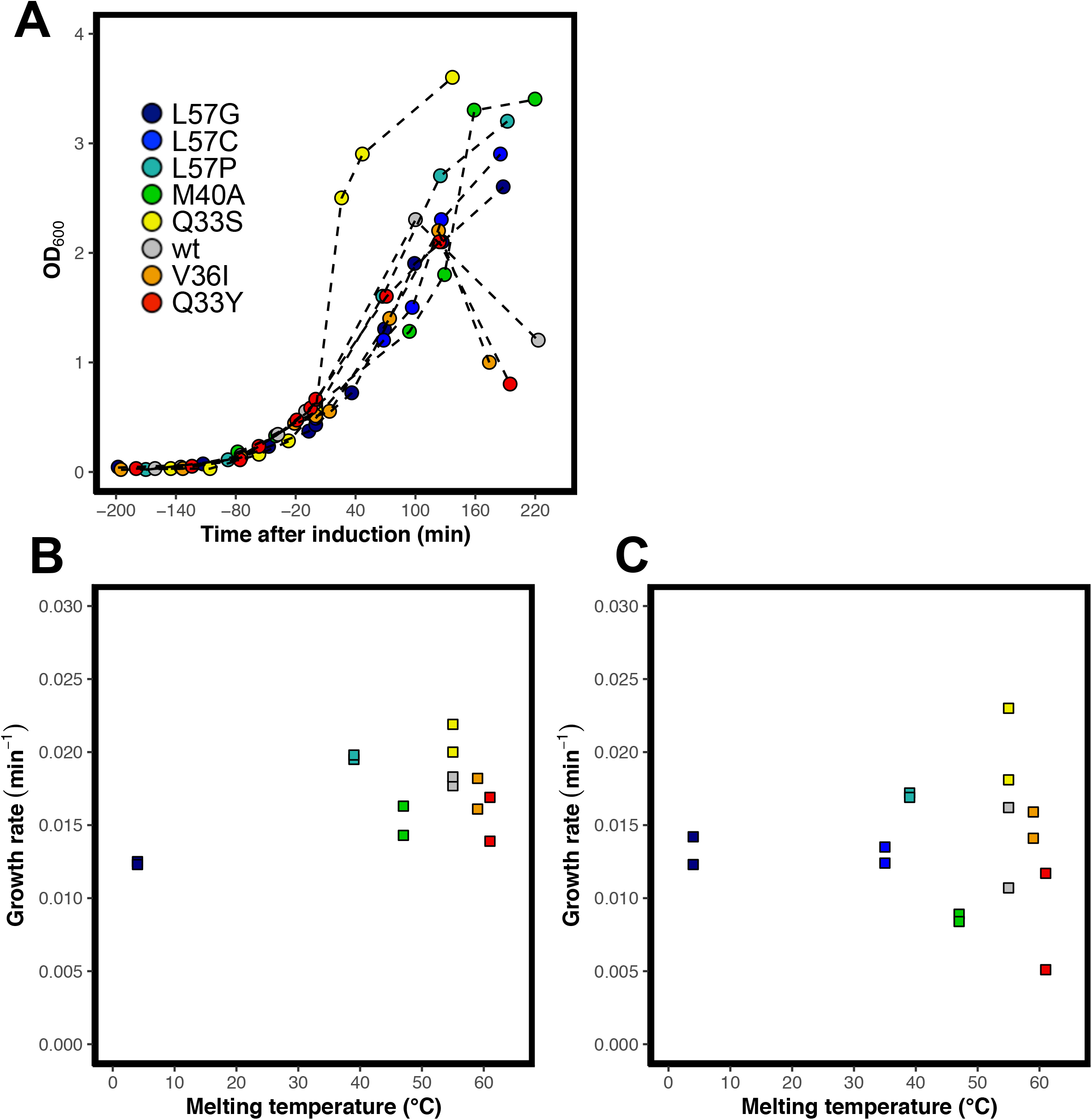
Growth of cells expressing N102LT variants. **A)** OD600 as a function of time after induction of protein synthesis for cells expressing the variants: L57G (dark blue), L57C (blue), L57P (cyan), M40A (green), Q33S (yellow), V36I (orange), Q33Y (red) and wt (grey). One of two replicates are shown for each variant. **B)** Growth rates prior to induction calculated for two replicates of each of the eight different N102LT variants with different thermal melting temperatures. Colour scheme as in 2A. Spearman rank correlation = 0.14. **C)** Growth rate the first 120 min after induction calculated for two replicates of each of the eight N102LT-variants with different melting temperature. Colour scheme as in 2A. Spearman rank correlation = 0.11.

### Cell growth after induction of recombinant protein production is variant-dependent

If variants trigger the PQC network to a varying degree, they might also have different impact on cell fitness. To investigate this, we tracked cell density during protein expression of each variant. We observe a small variant-dependent difference in growth rate even before induction of recombinant protein expression (Figure 2B), which is likely due to leaky expression. Independent of which variant is expressed, the cells continue to grow for approximately two hours after induction. After this point cells expressing the wt as well as Q33Y and V36I show a drop in OD_600_ (Figure 2A, S2) indicating that cells expressing these variants are dying. After induction of recombinant protein production there is greater variation in the growth rate of cells expressing different variants, but with similar patterns as before induction. Neither the preinduction nor postinduction growth rate correlates with protein stability (2B+C).

### Accumulation of protein is stability-dependent

To monitor the abundance of protein in the cell we followed the development of red fluorescence over time. The first 30 min after induction with IPTG the red fluorescence intensity is close to constant followed by continuous increase throughout the four hours for which protein production is monitored (Figure 3A + S3). Due to the long maturation time of tagRFP (t_50_∼42 min [35]) the low red fluorescence levels the first 30 min does not necessarily indicate a lack of protein synthesis. Throughout the entire experiment the red fluorescence correlates well with variant melting temperature (Figure 3B), with a spearman rank correlation of 0.83 after 120 min. At that point, variants with melting temperatures higher than the wt have red fluorescence intensities double to the variants with lower stabilities, with the exception of one replicate of Q33Y (Figure 3C). Looking at different mutations at the same site we observe that L57G and L57C have similar fluorescence, while the more stable L57P variant has an increased value. This indicates that the decreased expression in the former variants is due to stability rather than a consequence of mutating a particular position.

**Figure 3.**
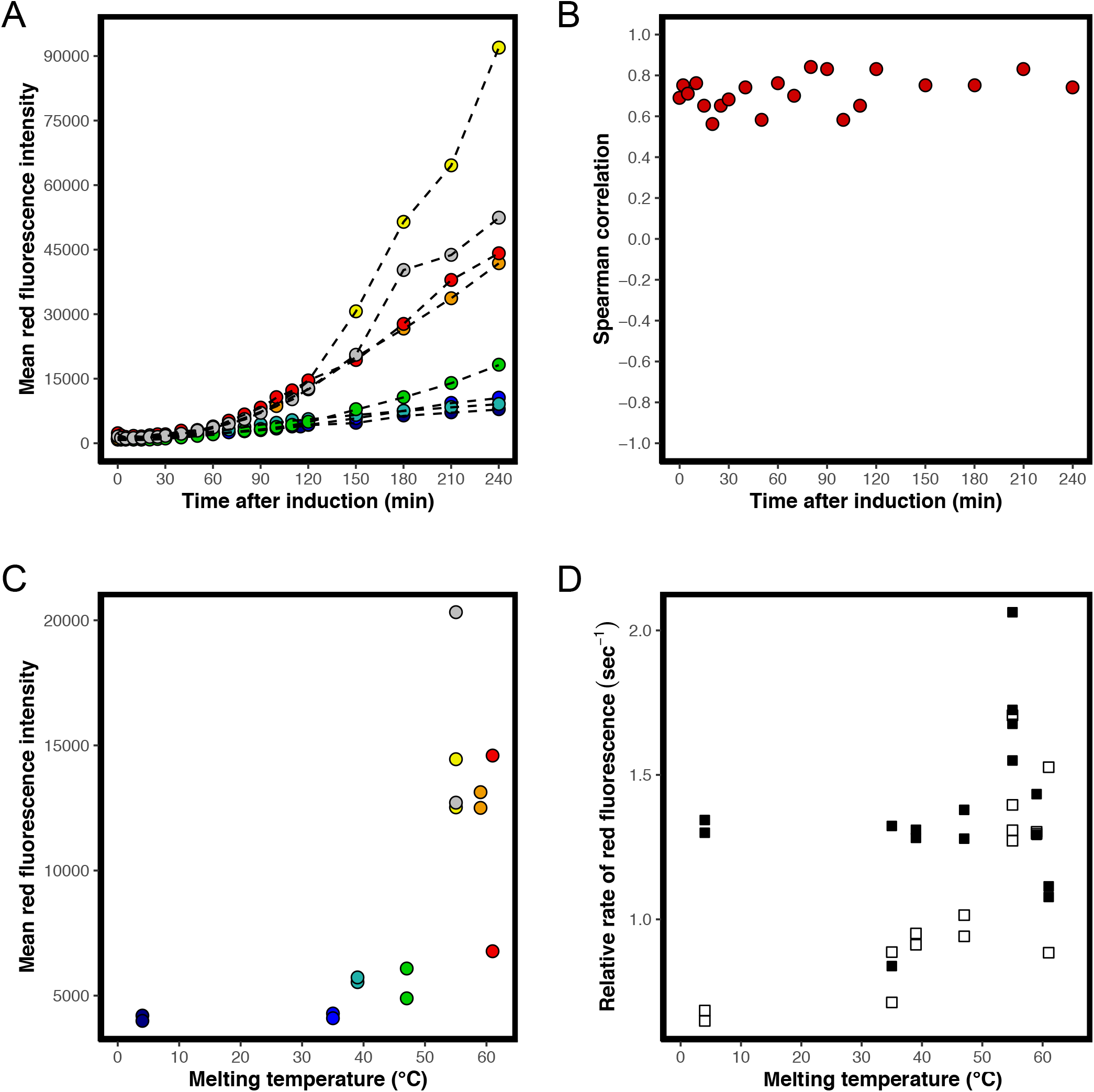
Abundance of tagRFP fusion proteins monitored by red fluorescence. **A)** Mean red fluorescence intensity of cells expressing variants: L57G (dark blue), L57C (blue), L57P (cyan), M40A (green), Q33S (yellow), V36I (orange), Q33Y (red) and wt (grey). One of two replicates are shown for each variant. **B)** Spearman correlation between mean red fluorescence and variant melting temperature at each time point. Calculations are based on two replicates of each variant. Colour scheme as in 3A. **C)** Mean red fluorescence intensity at the 120 min time point for the eight N102LT variants with different melting temperature. Spearman correlation = 0.83. Colour scheme as in 3A. **D)** Early (30 – 60 min, black) and late (60-90 min, white) relative rates of red fluorescence calculated for the eight N102LT variants with different melting temperature. Spearman correlation between early rate and variant melting temperature is 0.05. Spearman correlation between late rates and variant melting temperature is 0.70.

### Variation in protein accumulation is due to differences in rate of protein degradation rather than synthesis

The amount of accumulated protein is a consequence of the opposing effects of synthesis and degradation. Variation in abundance between mutants can thus be explained by differences in protein synthesis rate, protein degradation rate or both. We estimated the relative rate of red fluorescence change (change in fluorescence intensity divided by the intensity) between 30 and 60 min (“early” rates), and between 60 and 90 min (“late” rates) after induction. Both late and early rates varies with variant (Figure 3D). Interestingly, for variants less stable than wt the late rates are significantly reduced compared to the early rates. Assuming degradation is minimal early after onset of protein synthesis, this indicates that eventually the rate of degradation surpasses the rate of synthesis for these variants. For the more stable variants (wt, Q33S, Q33Y and V36I) the relative rate is not reduced or only minimally reduced after 60 min, which indicates that less protein is degraded in the cells harbouring these variants. The late rates correlate with thermostability with a spearman rank correlation of 0.7 (Figure 3D). Taken together, our results suggests that the difference in relative rate is most likely due to stability-dependent degradation of variants rather than difference in the rate of translation.

### Expression from the DnaK promoter is impacted by protein translation rates and thermostability

Protein folding stress triggers the PQC network and we wished to investigate whether protein translation alone is able to trigger a response and to what extent the biophysical properties, such as thermostability, of a protein impacts the stress response.

Assuming that the degradation of GFP is independent of the variant expressed, the total green fluorescence signal in Figure 4A and S4 reports on the total synthesis of GFP initiated at the stress-controlled DnaK promoter. Therefore, it can serve as a proxy for the accumulated DnaK response triggered by expression of the 8 variants. Immediately upon induction with IPTG an increase in green fluorescence is observed (Figure 4A and S4) which shows that the sfGFP signal emerge even before protein synthesis can be detected as an increase in red fluorescence. However, this can be explained by a shorter maturation time for sfGFP than tagRFP (14 min vs 42 minutes) [35]. Due to the much slower maturation time of tagRFP compared to sfGFP, total fluorescent intensities are not synchronized like the molecular events in the cell. However, by monitoring the rate of green and/or red fluorescence instead of total fluorescence we eliminate the effect of different maturation times.

**Figure 4.**
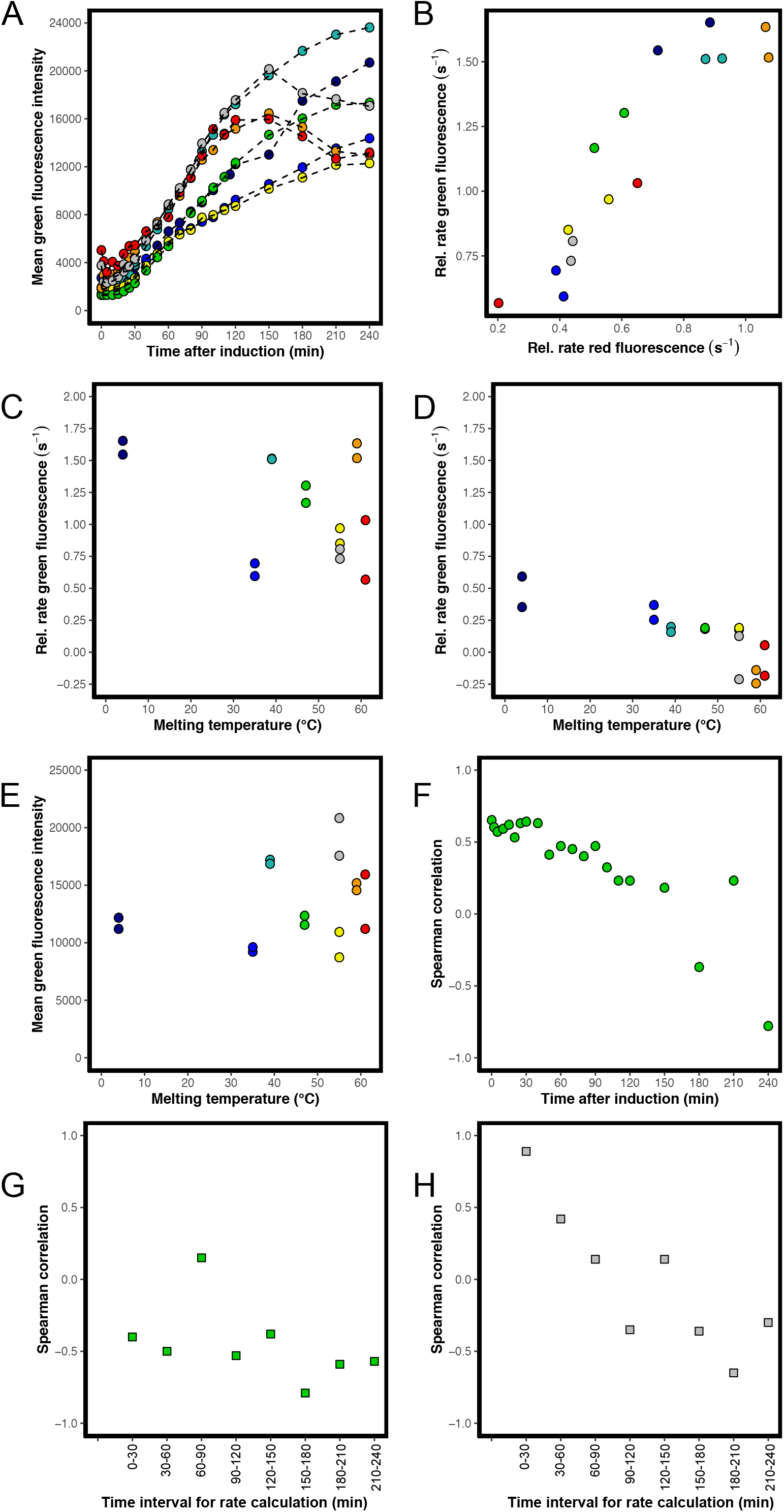
Transcription from the DnaK promoter monitored by green fluorescence. **A)** Mean green fluorescence intensity of cells expressing variants: L57G (dark blue), L57C (blue), L57P (cyan), M40A (green), Q33S (yellow), V36I (orange), Q33Y (red) and wt (grey). One of two replicates are shown for each variant. **B)** The initial (0-30 min) relative rate of green fluorescence as a function of the initial rate of red fluorescence. Spearman correlation = 0.89. Both replicates shown. Colour scheme as in 4A **C)** The initial (0-30 min) relative rate of green fluorescence calculated for the eight N102LT variants with different melting temperature. Spearman correlation between early rate and variant melting temperature is 0.40. Colour scheme as in 4A. **D)** The relative rate (150 min to 180 min) of green fluorescence calculated for the eight N102LT variants with different melting temperature. Spearman correlation between rate and variant melting temperature is 0.79. Colour scheme as in 4A. **E)** Mean green fluorescence intensity at the 120 min time point for the eight N102LT variants with different melting temperature. Spearman correlation between green fluorescence intensity and variant melting temperature is 0.23. Colour scheme as in 4A. **F)** Spearman correlation between accumulated mean green fluorescence and variant melting temperature at each time point. Correlations are calculated based on both replicates of each variant. **G)** Spearman correlation between rate of green fluorescence in the indicated time interval and variant melting temperature. Correlations are calculated based on both replicates of each variant. **H)** Spearman correlation between rate of green fluorescence and rate of rate fluorescence in the indicated time interval. Correlations are calculated based on both replicates of each variant.

#### Protein translation alone triggers DnaK transcription

The initial rate (0 to 30 min after IPTG induction) of green fluorescence correlates strongly with the initial rate of red fluorescence (spearman rank correlation of 0.89) (figure 4B), but poorly with the melting temperature of variants (Figure 4C). This observation suggests that at the onset of protein synthesis the DnaK promoter is upregulated purely in response to increased protein translation – independent of the stability of the expressed protein. Later in the time course the rate of red fluorescence no longer corresponds to the rate of translation due to increased influence of degradation, and the correlation between the relative rate of red and relative rate of green fluorescence diminishes (Figure 4H).

#### Variants with low stability show increased DnaK transcription

In contrast to what is observed at early rates, there is a moderate correlation between rate of GFP and melting temperature following the first 60 min of protein synthesis. The correlation appears to increase at later timepoints (Figure 4D and 4G). While individual correlation values fluctuate and are associated with statistical uncertainty, there is a clear indication that the DnaK response is affected by the thermal stability of the variants (Figure 4G). Notably, after 120 min, cells expressing the most stable variants (wt, Q33Y and V36I) experience a significant drop in green fluorescence (Figure 4A+D). This could suggest that the stable variants do not engage DnaK, likely due to lower concentration of unfolded/misfolded proteins. However, concurrently with a drop of in green fluorescence cells expressing these three variants also experience a decrease in OD. Consequently, the decrease in expression from the DnaK promoter could be a trait associated with cell death. The green fluorescence in cells expressing the remaining five variants continue to increase, although with reduced rates (Figure 4A+D) and with higher rate of green fluorescence for the less stable variants. A correlation between the rate of green fluorescence and melting temperature is observed even when considering only the subset of five with positive GFP rates, but only after 150 minutes (spearman rank correlation 0.78 for rates calculated in the range 180-210 minutes and 0.64 for the 210-240 interval). So, as could be expected, our assay indicates an increased requirement for chaperone assistance for the less stable variants.

The first 50 min post-induction there is a moderate correlation between melting temperature and accumulated green fluorescence. However, onwards from 60 min the correlation decreases steadily (4F). This most likely reflects the variation in protein abundance as the effect of degradation of the N102LT variants becomes increasingly pronounced. To study how much the expression and stress response vary between individual cells, we must consider differences in plasmid copy number in cells. Since both RFP and GFP are expressed from the same plasmid we can account for this by evaluating the ratio between GFP and RFP signal. This ratio also provides a metric of the chaperone response relative to protein abundance. For all variants we see an increase in green to red fluorescence ratio up until around 30 minutes after induction (Figure 5A, S5). After that point, the normalised DnaK response decreases in a manner highly dependent on the expressed variant (Figure 5B). In addition, the accumulated and normalised DnaK response correlates strongly with the melting temperature of the variant following the initial peak at 30 min and throughout the entire 240 min (Figure 5D). This is exemplified by the 120-minute time point, where we see a clear distinction between the chaperone response triggered for the eight N102LT variants, with a strong negative correlation between the green to red fluorescence ratio and the melting temperature of the expressed variants (Figure 5C). Less stable variants induce a stronger DnaK response, and that pattern persists over time. This observation is a strong indication that the less stable variants require continuous assistance from PQC chaperones to maintain their native fold, whereas more stable variants do not.

**Figure 5.**
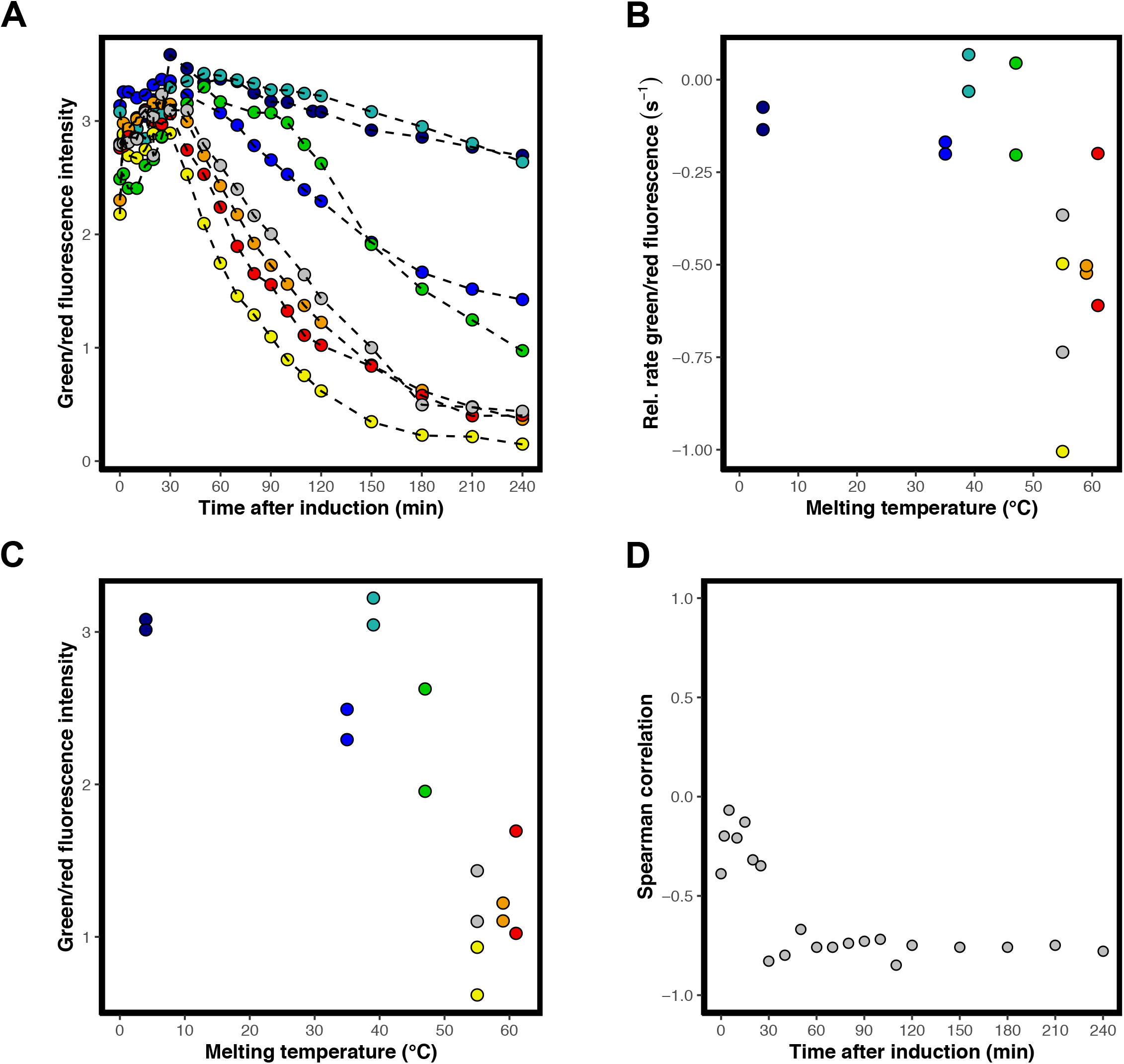
Accumulated transcription from the DnaK promoter normalised to protein abundance. **A)** Mean green fluorescence intensity divided by red fluorescence intensity of cells expressing variants: L57G (dark blue), L57C (blue), L57P (cyan), M40A (green), Q33S (yellow), V36I (orange), Q33Y (red) and wt (grey). One of two replicates are shown for each variant. **B)** Relative rate of green/red fluorescence (30-60 min) as a function of melting temperature. Spearman correlation = 0.55). Colour scheme as in 5A **C)** Mean green fluorescence intensity divided by red fluorescence intensity at the 120 min time point for the eight N102LT variants with different melting temperature. Spearman correlation between green/red fluorescence intensity and variant melting temperature is 0.75. Colour scheme as in 5A **D)** Spearman correlation between green/red fluorescence intensity and variant melting temperature at each time point. Calculations are based on both replicates of each variant.

### Expression of the wt, Q33Y and V36I variants leads to occurrence of subpopulations with altered fluorescence phenotypes

The above analysis shows that evaluation of protein abundance as well as DnaK response gives a good insight into the effect of protein biophysical properties on whole populations of cells expressing the same variant. However, with flow cytometry we can go beyond the study of cells at the population level and investigate the cell-to-cell variation in protein expression and DnaK response. To fully explore the phenotypic variation within a population, we cluster the cells based on red and green fluorescence using a Bayesian Gaussian mixture model. Although not all identified clusters may represent homogenous phenotypes, automated clustering allows for unbiased assignments. Figure 6 shows the clusters identified for variant L57G and V36I at times 0, 90, and 240 min in a double logarithmic plot. If the cells form a straight line with a slope of one in the double logarithmic plot the green to red fluorescence ratio is constant in all cells and is equivalent to the intersection with the y-axis. A slope below one indicates that the variation in red fluorescence is larger than the variation in green fluorescence, whereas a slope above one indicates that the variation in green fluorescence is bigger than the variation in red fluorescence.

**Figure 6.**
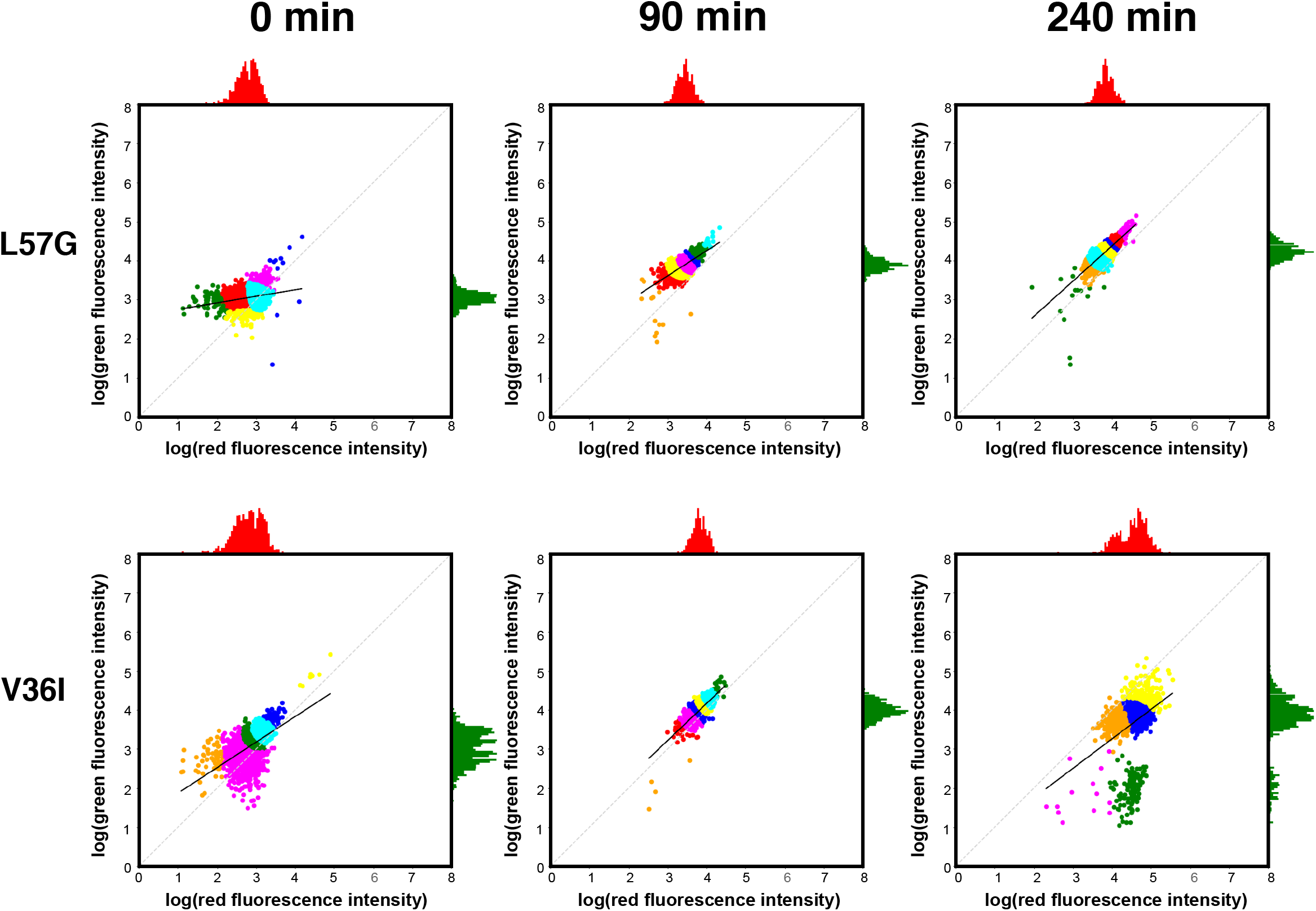
Red and green fluorescence of single cells expressing L57G and V36I. The red and green fluorescence for each individual cell in a population expressing L57G (top) and V36I (bottom) at the time of induction (left) and 90 min (middle) and 240 min (right) after induction with IPTG. Colouring in scatter plot is based on a Bayesian Gaussian mixture model for cluster assignment. The x-axis histogram (red) shows the distribution of red fluorescence, and the y-axis histogram (green) shows the distribution of green fluorescence within the population.

For all variants we identify one main population consisting of 3 to 4 different clusters (Figure 6, S6-7). The distribution of red and green fluorescence within the main population differs between variants and with time. At the time of induction, both green, and especially red fluorescence, show significant cell-to-cell variation for all variants. This is evident by the dispersed distribution around a line with a slope below one and is observed regardless of variant. However, for some variants there is an indication of a diagonal line pre-induction. This is likely due to leaky expression. Once recombinant protein synthesis is induced, the distribution of red and green fluorescence in the main population becomes increasingly homogenous and approaches a near-constant ratio between green and red fluorescence for all cells in a population (Figure 6, Figure 7, S6-8). Interestingly, when expressing variants with a melting temperature below the wt, reaching the near-constant ratio is significantly delayed (Figure 7, S8).

**Figure 7.**
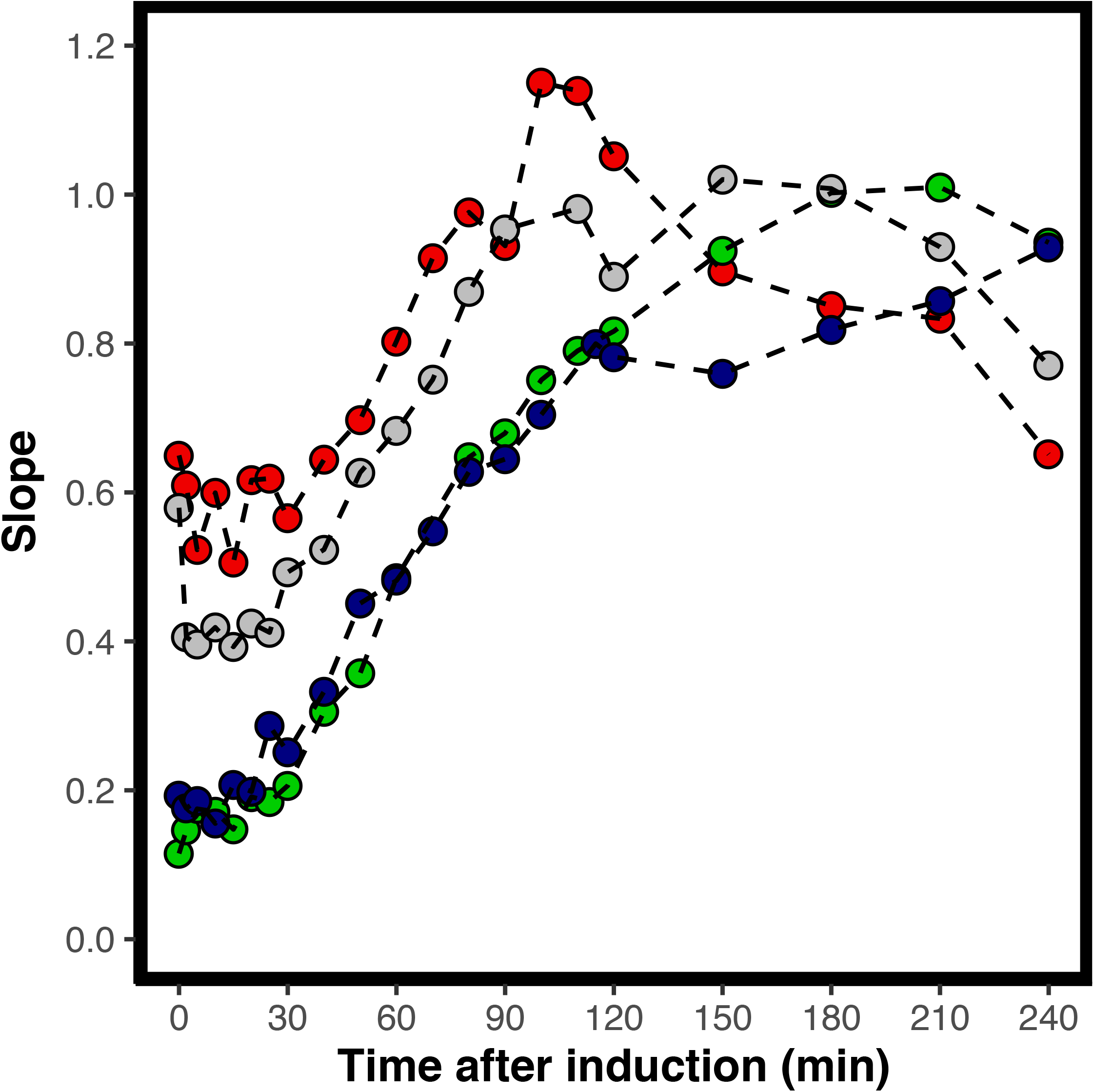
Slope of fluorescent population over time. For each time point the slope of log transformed red fluorescence vs log transformed green fluorescence is calculated for L57G (dark blue) M40A (green), Q33Y (red) and wt (grey). One of two replicates are shown for each variant.

For the least stable variants (L57G, L57C, L57P and M40A) and Q33S, with a stability close to wt, there appears to be only a single population (Figure 6+S6-7). For wt and the two most stable variants (Q33Y and V36I), however, individual cells with less protein abundance (lower red fluorescence intensity) emerge after 120 min, and the fraction of these cells increase over time. After 240 minutes, distinct populations with phenotypic variation in DnaK response can be observed for cells expressing wt, Q33Y and V36I. Cells with low green fluorescence have either low or high red fluorescence, forming two distinct populations. We note that the emergence of subpopulations with altered fluorescence phenotypes are highly variant-specific and is only observed for cells expressing the three most stable variants and concur with a reduction in OD for cells expressing these variants. Neither formation of subpopulations nor reduction in OD is observed for cells expressing Q33S which has thermostability equivalent to the wt.

### Morphological changes are observed for E. coli expressing N102LT variants

Stressed cells often show morphological changes. We monitored cell morphology by forward (FSC) and side scatter (SSC) to identify potential variant-dependent changes. FSC provides information about the size of the cell, and for most variants there is no change in FSC over the four-hour time-course indicating that the size of the cells remains constant (Figure 8A, S9). However, for three variants (wt, Q33Y and V36I) the FSC increases after 120 minutes, indicating an increase in cell size. Furthermore, these three variants are distinctly different from the other five variants throughout the timeseries, indicating that cells expressing wt, Q33Y and V36I might be bigger even prior to induction. For eukaryotic cells SSC reports on the complexity of the cell but it remains unclear how to interpret SSC from bacterial cells, although it has been suggested that SSC is associated with cell wall density [36]. We observe a variant-dependent decrease in SSC during the first two hours after induction Figure 8B, S10) with a moderate negative correlation to the melting temperature of the expressed variant. Although the implications of SSC are not understood it is evident from the patterns of FSC and SSC variation that cells undergo morphological changes, and that the changes depend on the expressed variant.

**Figure 8.**
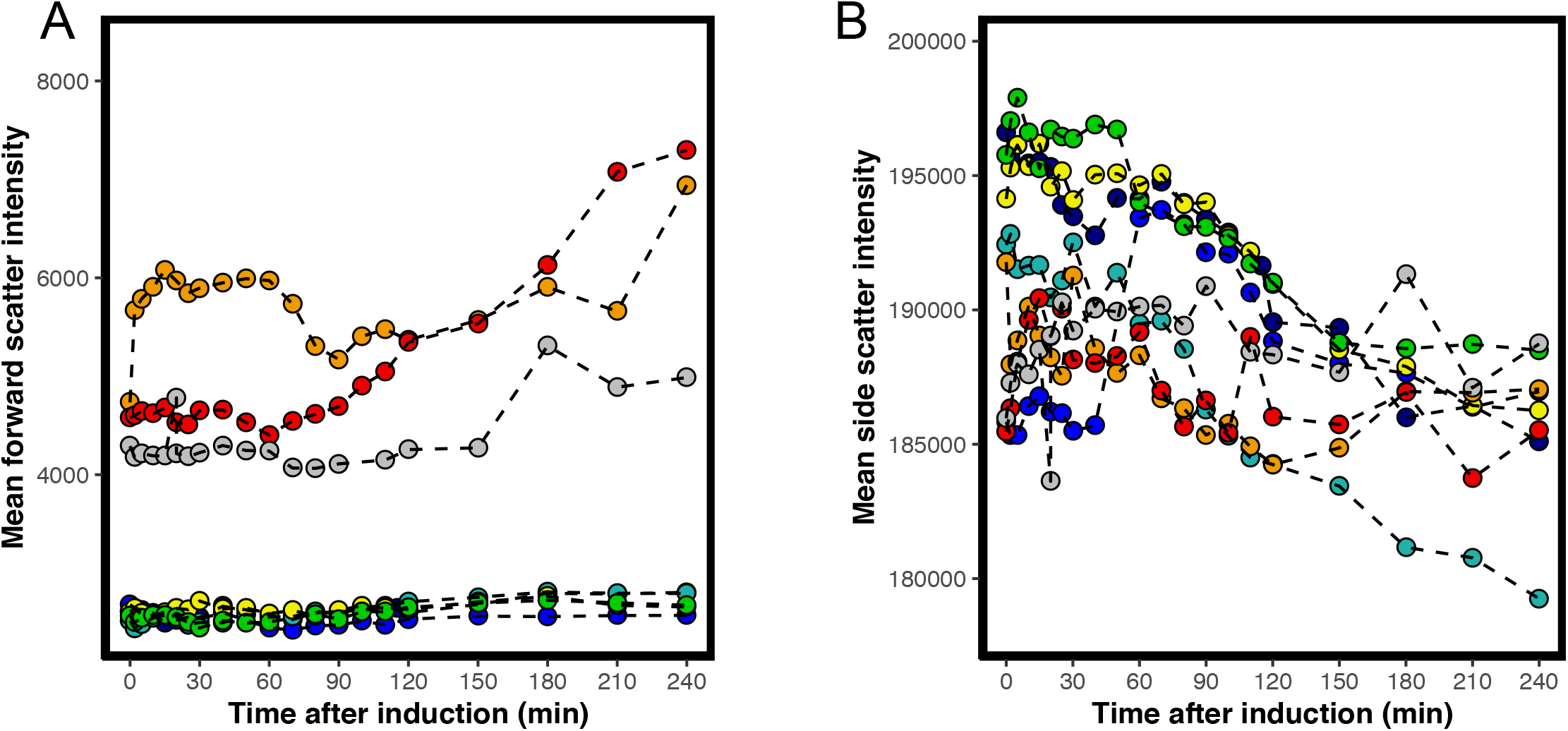
Cell morphology of whole populations expressing N102LT variants. Forward (**A**) and side (**B**) scatter as a function of time for variants: L57G (dark blue), L57C (blue), L57P (cyan), M40A (green), Q33S (yellow), V36I (orange), Q33Y (red) and wt (grey). One of two replicates are shown for each variant.

## Discussion

### The DnaK response is triggered by protein translation rate during protein synthesis and thermostability at folding equilibrium

We have demonstrated that DnaK expression is activated immediately once recombinant protein expression starts. It then shifts from being dependent on protein translation rates to being primarily governed by protein stability at later time points. The initial, stability-independent activation is most likely due to the increase of unfolded protein as the nascent peptide chain exits the ribosome. The observed correlation between onset of protein synthesis and initial DnaK response can be caused by either co-translational binding of DnaK to the emerging peptide chain or binding of DnaK to newly synthesised, unfolded, protein after it is released from the ribosome (Figure 9). However, several *λ*-repressor constructs have been shown to fold in submilisecond time scales [37-39] why presence of significant amounts of unfolded *λ*-repressor are unlikely. Binding of DnaK to the unfolded protein releases DnaK- bound *σ*^32^ which acts as transcription factor for synthesis of DnaK as well as other PQC proteins. The consequence is an increase in transcription of PQC proteins that are unrelated to protein thermostability but tightly linked to translation rate. Besides aiding in nascent chain folding, this linkage adjusts the amount of PQC network components to the overall protein expression level and thus ensures that enough chaperones are present to deal with potential protein misfolding. The rate of GFP synthesis as well as accumulated GFP/RFP signal correlates with variant melting temperature after the initial 30 minutes of protein synthesis. This correlation suggests that the DnaK response is impacted by the proportion of protein in the unfolded/misfolded state at folding equilibrium (Figure 9). Hence, variants with lower stability require increased DnaK activity compared to more stable variants. Increased interaction with DnaK increases the degree of degradation by proteases, as DnaK, given its role as a central hub in the PQC network, is involved in targeting misfolded proteins for degradation [40]. As a consequence, in our study, most protein is accumulated in cells expressing the four variants that are at least as stable as the wt (wt, Q33S, Q33Y and V36I), and thus does not require continuous assistance from chaperones.

**Figure 9.**
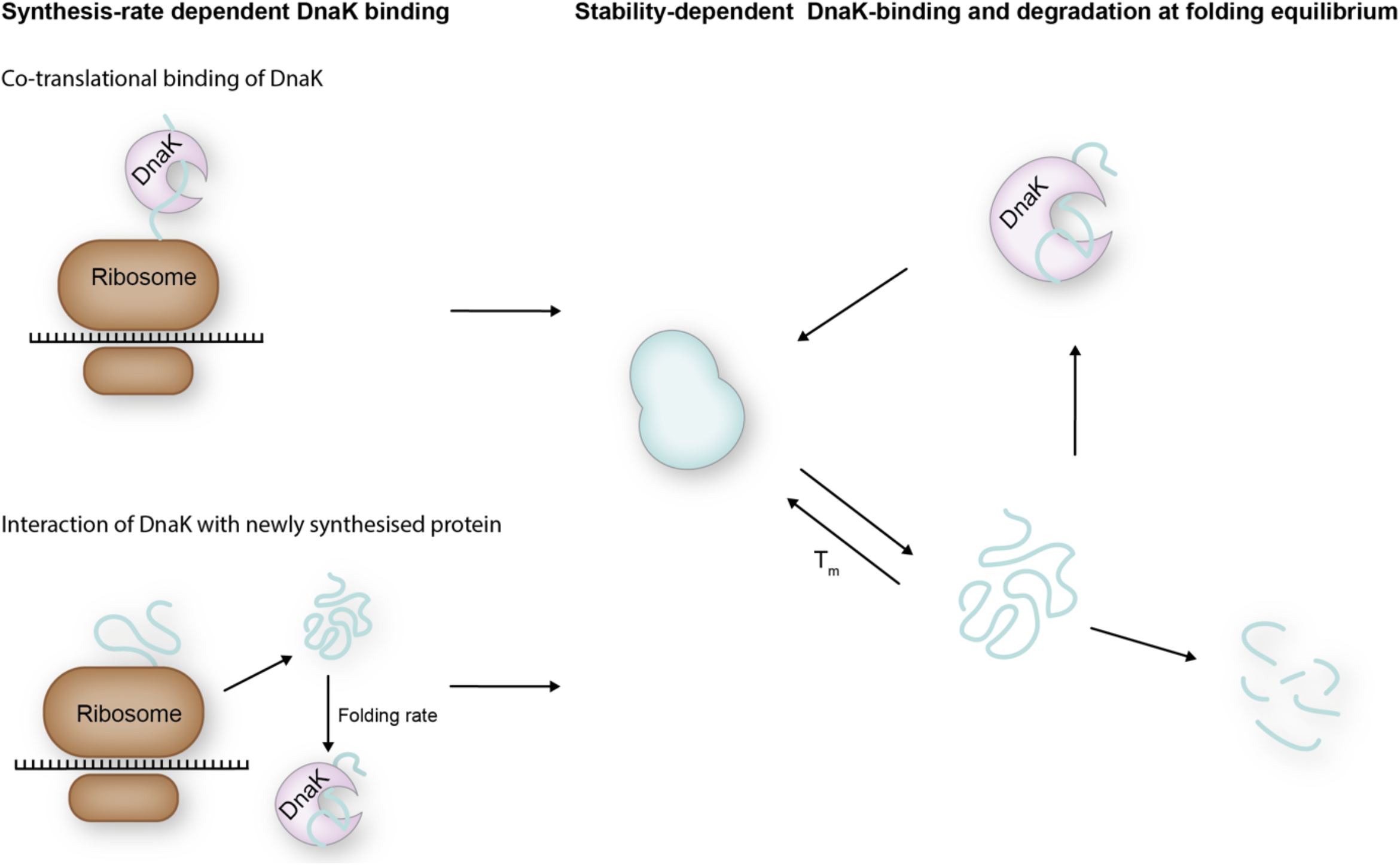
A model for DnaK interaction with client proteins. Two different factors impact the interaction of DnaK with client proteins: synthesis rate and protein stability. DnaK can act co-translationally with the nascent peptide chain as it exits the ribosome exit tunnel, in which case DnaK response is expected to be proportional to the translational rate. In addition, DnaK can bind to unfolded newly synthesised protein that has been released from the ribosome. For slow folding protein the interaction with DnaK will then also depend on folding rate. Fast folding protein will interact only minimally with DnaK. Fully synthesised proteins are in an equilibrium between folded and misfolded determined by the stability of the protein. Thus, DnaK binding to misfolded protein at folding equilibrium is dependent on protein stability

### Increased cell-cell variation at late time points for stable variants

Analysis of the accumulated DnaK response and N102LT variant accumulation in single cells revealed that cell-cell variation, especially in protein accumulation, is bigger early after onset of protein synthesis and decreases with time. There is a prolonged period with a wide distribution in protein accumulation observed for variants less stable than wt. This lag can potentially be explained by increased degradation: The variation in protein abundance can be caused by a variance in both protein synthesis and degradation. The variance for the more stable variants, however, is primarily due to variation in synthesis. Once folding and synthesis/degradation equilibrium is reached, a single population of cells is seen, with similar chaperone response relative to protein abundance. This suggests that as the cells reach some sort of homeostasis, the variation in DnaK response can be largely explained by variation in accumulated protein.

However, for the wt, Q33Y and V36I variants subpopulations with reduced DnaK response emerges after approximately 120 minutes. In association with the emergence of subpopulations in these three stable variants there is a decrease in OD_600_ and DnaK as well as an increase in cell size (a phenotype often associated with stressed cells [41]) is observed. The transformation into these alternative phenotypes is potentially related to the burden associated with a high protein load in the cell. Despite the Q33S reaching the highest overall abundance of protein, this variant does not share the described phenotype indicating a more complex relationship between protein stability and cellular response.

### Sequence-dependent effect on *in vivo* folding equilibrium or solubility might explain effects not accounted for by thermostability

Although we find a good overall correlation between the variant melting temperature and the DnaK response after expression in *E. coli*, not all the variation can be explained by the altered thermostability. Consequently, there might be other sequence-dependent effects that we have not investigated further. The folding equilibrium might be shifted *in vivo* compared to the conditions *in vitro*. The unfolded state for instance, could be stabilized by inadvertent interactions with other macromolecules. Similarly, the folded state could be further stabilised by non-specific interactions, e.g. with DNA. Altered affinity for DNA could also affect a variant’s toxicity and potentially explain the altered phenotypes observed exclusively for the wt, Q33Y and V36I which cannot be explained by thermostability alone. The DnaK-dependent signal is further potentially skewed by a variant’s aggregation propensity as aggregation itself has an impact on PQC network activity [14].

### Differential PQC response could constrain evolution of protein sequences

Several studies have investigated the coupling between biophysical properties and protein evolution. A strong negative correlation between rates of protein coding sequence evolution and gene expression levels have been observed [11, 42]. This has been coupled to protein stability by two different models. In the first, unstable proteins have a lower concentration of active protein resulting in lower fitness. In the second, the misfolded proteins are toxic to the cell with an associated fitness cost. Mathematically, these models become identical for proteins with stabilities beyond 5 kcal/mol [43]. Geiler-Samerotte et al. [14] directly tested the misfolding hypothesis by evaluating growth rates in yeast expressing misfolded proteins. They found a significant fitness cost due to misfolded proteins and associated upregulation of HSP70 (DnaK). The increased evolutionary rate of proteins with stronger binding to DnaK suggests that chaperones can buffer deleterious mutations [10]. Nonetheless, triggering the PQC network comes with a cost and it is possible that this burden on the cell can impact selection of protein variants during evolution. The observation that small amino acid changes can have a significant impact on the chaperone response thus suggests that the degree of *dnaK*-activation by a protein may itself act as a significant selection factor. The exact relation between magnitude of the PQC and fitness, however, is not clear. In our experiments, we did not observe a correlation between DnaK response and cell growth rates, but such correlations are likely obscured by variations in protein abundance. Moreover, the phenotypic differences can be very small to still have an effect on evolutionary time-scales. It should be noted that we carry out our experiments under conditions of strong protein overexpression. Hence, it is not clear if our results are relevant to evolution of protein sequences under more standard conditions.

### The reporter plasmid has potential use for screening protein stability

Stability is often a prerequisite for the application of proteins in research, medicine and biotechnology. The development of accurate and high-throughput assays that can screen for stability independent on protein function are thus desired. Our data demonstrate strong correlations between *in vitro* protein stability and fluorescence values recorded from single cells expressing the N102LT variants. This highlights the utility of the fluorescent reporter plasmid in combination with flow cytometry analysis as an assay for screening for protein stability *in vivo*. We propose two potential approaches for using the reporter assay for screening: (i) using the GFP/RFP ratios of tagRFP fusion proteins as a high throughput screen for variant libraries, e.g. for directed evolution. The strong correlation between GFP/RFP ratio and thermostability provides optimal conditions for screening for variants with increased stability since the correlation varies minimally with time. (ii) using the rate of GFP for high-throughput stability screening of protein variants. This approach is potentially less robust than the GFP/RFP ratio method but has the advantage of not requiring the proteins of interest to be fused to tagRFP. This can for example be beneficial in cases where the studied protein is involved in multimerization reactions where the large tagRFP molecule can inhibit proper assembly.

## Material and methods

### Strains and plasmids

A modified pQlinkN plasmid [44] were created by inserting additional restriction sites for PstI, SacII and NdeI between the Link1-region and the multiple cloning site (MCS) of pQlinkN. The genes encoding sfGFP and the dnaK-promoter (P_DnaK_) was amplified by PCR from *E*.*coli* BL21*(De3) genomic DNA, respectively. Both *sfgfp* and *P*_*dnak*_ was inserted into the modified pQlinkN plasmid using restriction enzyme cloning.

The gene for N102LT N-terminally fused with tagRFP (gi: 336287738) was synthesised (GenScript) and inserted to unmodified pQlinkN plasmid using restriction enzyme cloning. Point mutations in the N102LT gene were made by site-directed mutagenesis with Phusion HotStart II DNA polymerase to create seven N102LT variants: Q33S, Q33Y, V36I, M40A, L57C, L57G and L57P. The full reporter plasmid was created by ligation independent cloning with pQLinkN harbouring *N102LT-tagRFP* variants as recipient vector and modified pQlinkN harbouring *P*_*DnaK*_ and *sf-gfp* as insert vector [44].

### Protein expression

*E. coli* BL21(DE3) pLysS was transformed with reporter plasmid harbouring the N102LT variants for expression. Cells were grown at 37 °C in LB medium with shaking (150 rpm) and were induced with 1 mM isopropyl-β-d-thiogalactopyranoside (IPTG) when cells reached an OD_600_ between 0.4 and 0.6 and continued to grow for four hours. Variants were expressed on separate days, but both replicates of the one variant was expressed individually on the same day.

### Flow cytometry

Culture samples were taken just before induction with IPTG (time point 0 min), after 2 min, every five min the first 30 min, every 10 min from 30 min to 120 min and every half hour until 240 min after induction. Immediately after taken the sample it is diluted diluted 1:1000 in 10 mM Na_2_HPO_4_, 1.8 mM KH_2_PO_4_, 137 mM NaCl, 2.7 mM KCl pH =7.4 and analysed on s3e cell sorter (Biorad). Replicates were measured immediately after each other giving a time lag between replicates of 10-30 sec. 3000 events were collected with an event rate of ∼500 events/s. A laser with 488 nm was used for detection. Cells expressing N102LT variants were gated on forward and side scatter to exclude doublets. Fluorescence was detected using a 525/30 filter (green fluorescence) and a 615/25 filter (red fluorescence). Flow cytometry data was analysed using the flowCore [45], flowViz [46] and flowWorkspace [47] packages within the Bioconductor frame work.

### Clustering

Flow cytometry data was clustered using Bayesian Gaussian mixture model, implemented in scikit-learn [48]. Up to 7 different components were allowed and priors for weights were described by a Dirichlet distribution with a weight concentration prior of 1e-6. A diagonal covariance matrix was used for each component.

### Data analysis

Mean red and green fluorescence as well as mean forward and side scatter intensities were calculated from the absolute intensity for each event for a given variant at a given time. Mean green/red fluorescence intensity was calculated by taking the mean over the green/red fluorescence intensity for each variant for each timepoint. Rates were estimated by fitting a linear trendline to the mean red, green or green/red fluorescence intensity within the indicated time interval. Relative rates were estimated by normalizing the rate of fluorescence with the mean fluorescence within the same time interval. Mean red, green and green/red fluorescence as well as relative rates were calculated individually for each replicate. Spearman correlations between red, green or green/red and variant melting temperature were calculated based on both replicates of each variant. A linear trend line was fitted to the log10 transformed red and green fluorescence for each event for a given variant at any given time point. A trend line was fitted individually for each replicate.

## Acknowledgements

We would like to thank Dr. Fabian Cornejo for helpful comments to the manuscript. We would also like to thank Robert T. Sauer for providing the original N102LT construct.

## Author contributions

Sebastian Rämisch: Methodology, Writing – Review and editing

Signe Christensen: Conceptualization, Methodology, Formal analysis, Investigation, Writing – original draft, Visualization

Ingemar André: Conceptualization, Methodology, Software, Resources, Supervision, Writing original draft

## Conflict of interest

The authors declare that they have no conflict of interest.

## Funding

This work was supported by the European Research Council (ERC) under the European Union’s Horizon 2020 research and innovation programme [771820]

---

**S 1.**
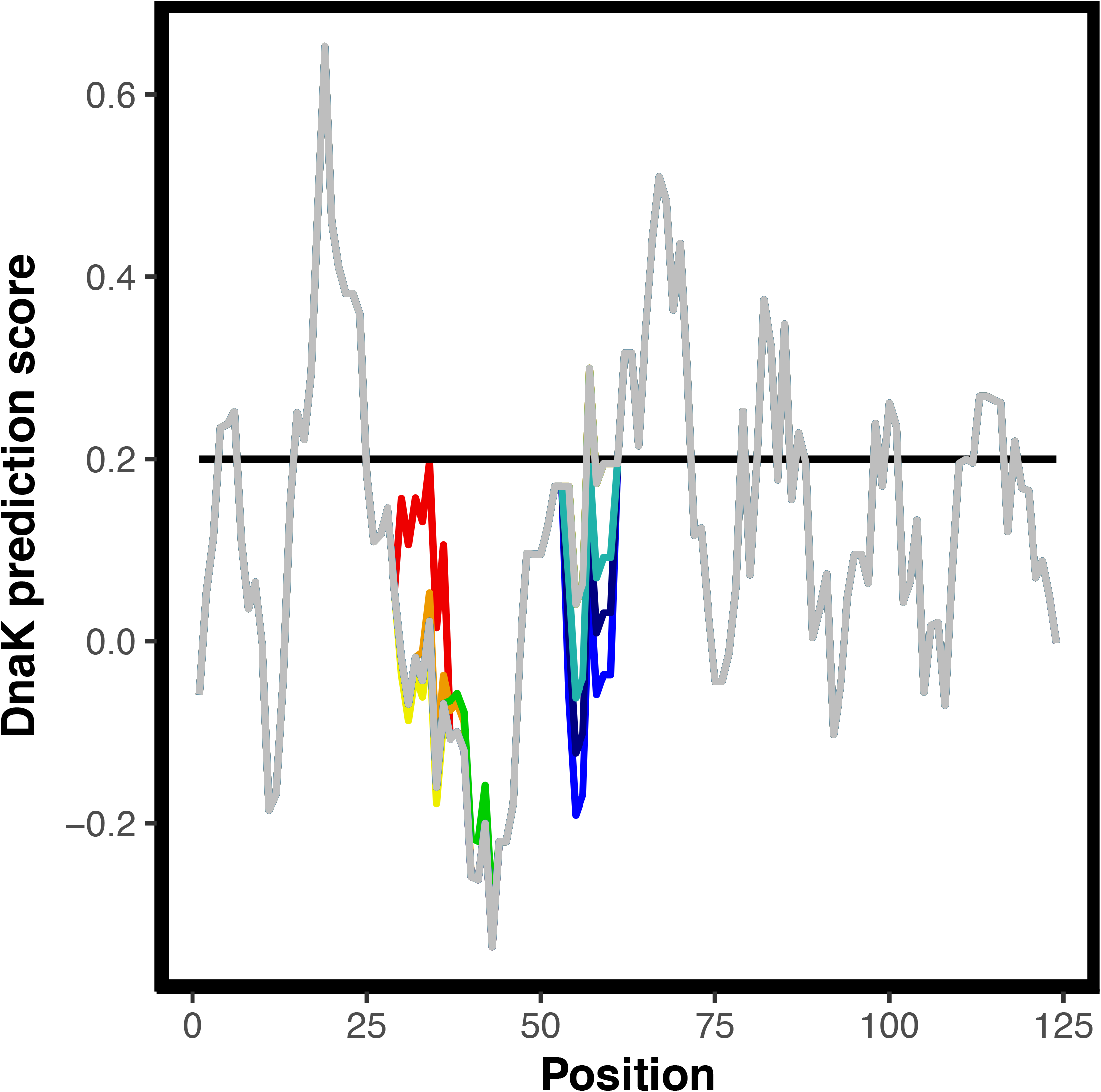
Prediction of DnaK binding sites in N102LT variants. DnaK binding sites predicted by the qualitative ChaperISM algorithm {Guiterres, 2020} for variants: L57G (dark blue), L57C (blue), L57P (cyan), M40A (green), Q33S (yellow), V36I (orange), Q33Y (red) and wt (grey). Where only grey is visible the score is identical for all variants. Cut-off for DnaK binding is 0.2 (black line).

**S 2.**
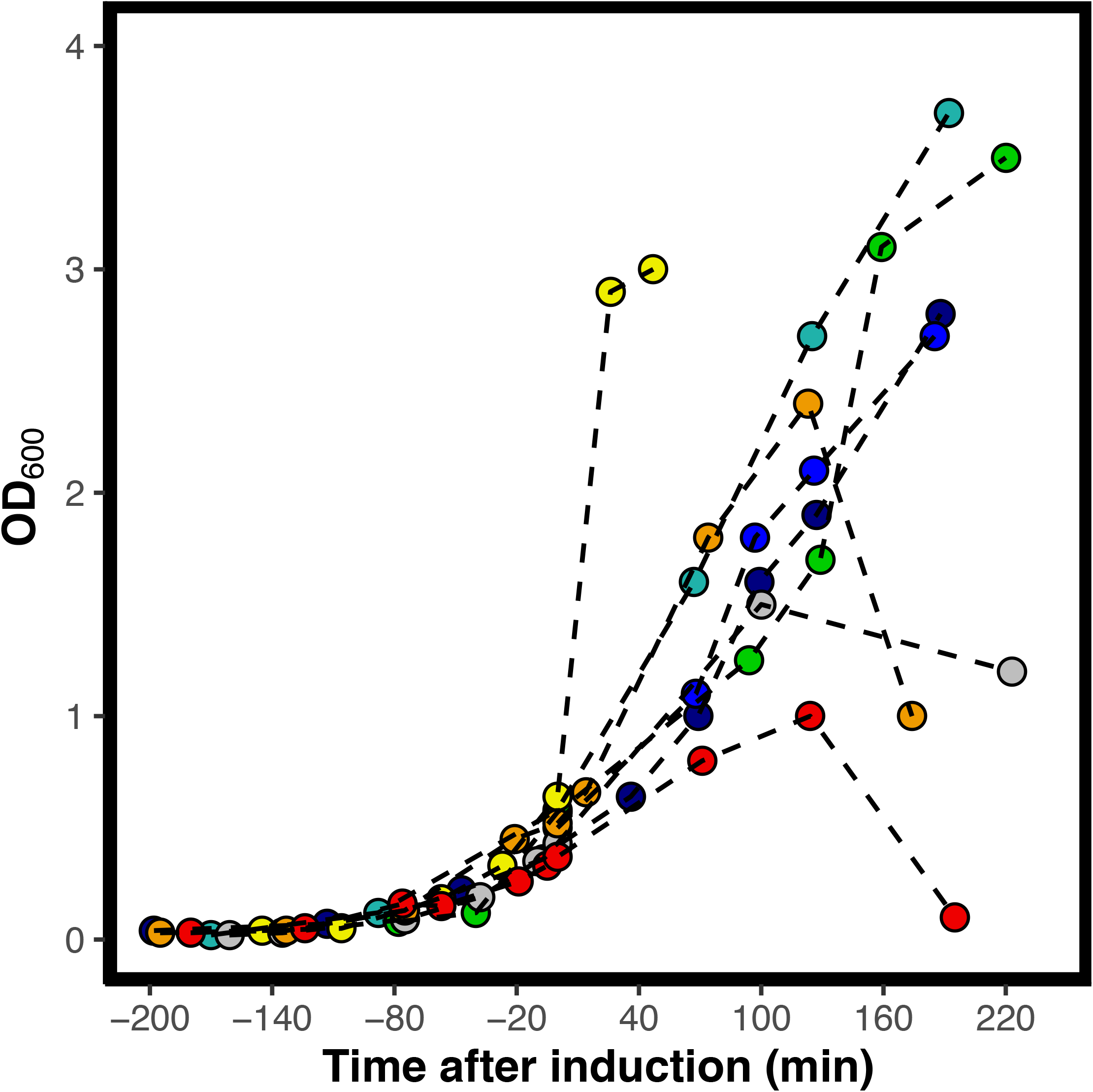
Growth of cells expressing N102LT variants. OD600 as a function of time after induction of protein synthesis for cells expressing the variants: L57G (dark blue), L57C (blue), L57P (cyan), M40A (green), Q33S (yellow), V36I (orange), Q33Y (red) and wt (grey). One of two replicates are shown for each variant.

**S 3.**
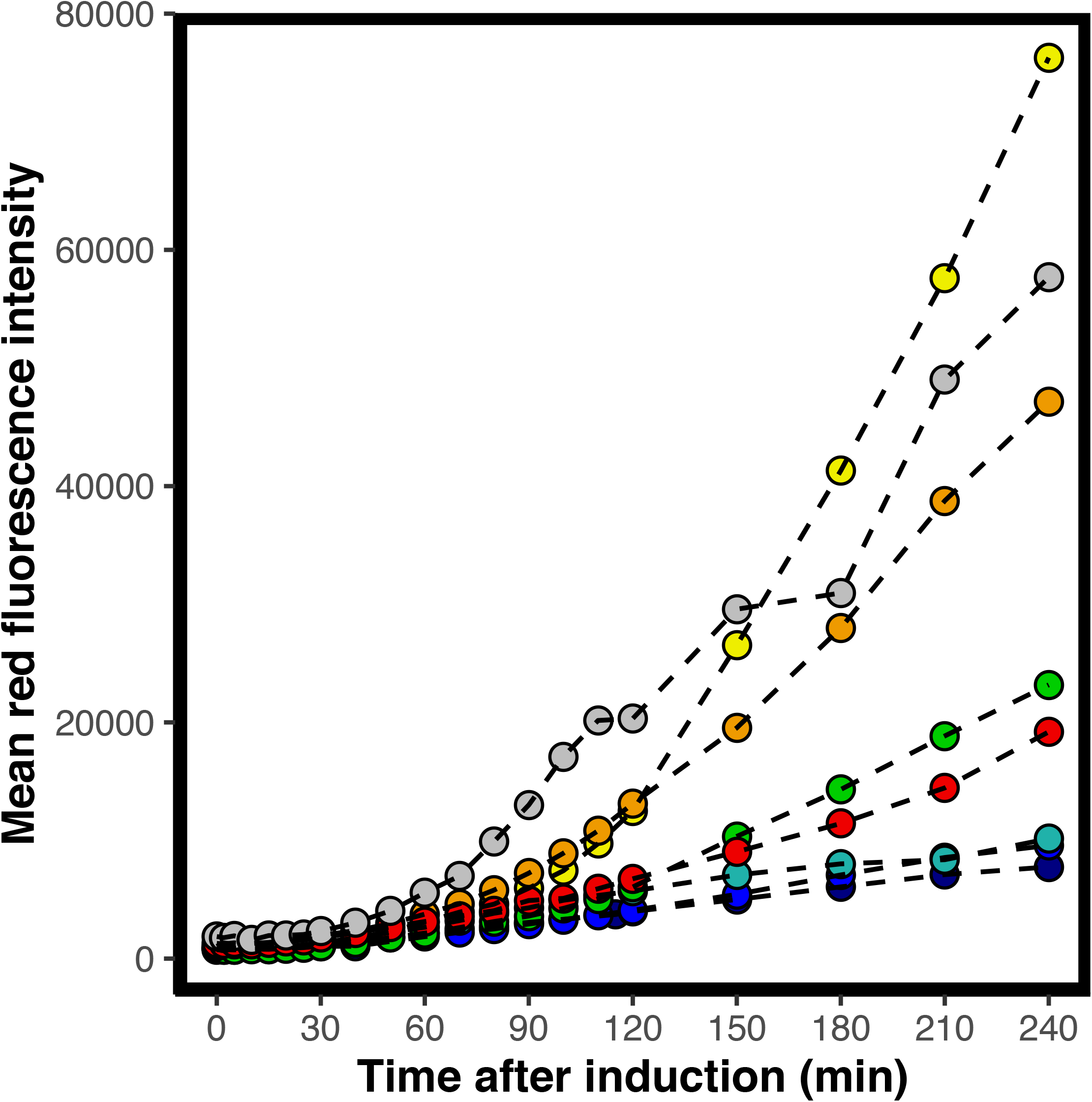
Abundance of tagRFP fusion proteins monitored by red fluorescence. Mean red fluorescence intensity of cells expressing variants: L57G (dark blue), L57C (blue), L57P (cyan), M40A (green), Q33S (yellow), V36I (orange), Q33Y (red) and wt (grey). One of two replicates are shown for each variant.

**S 4.**
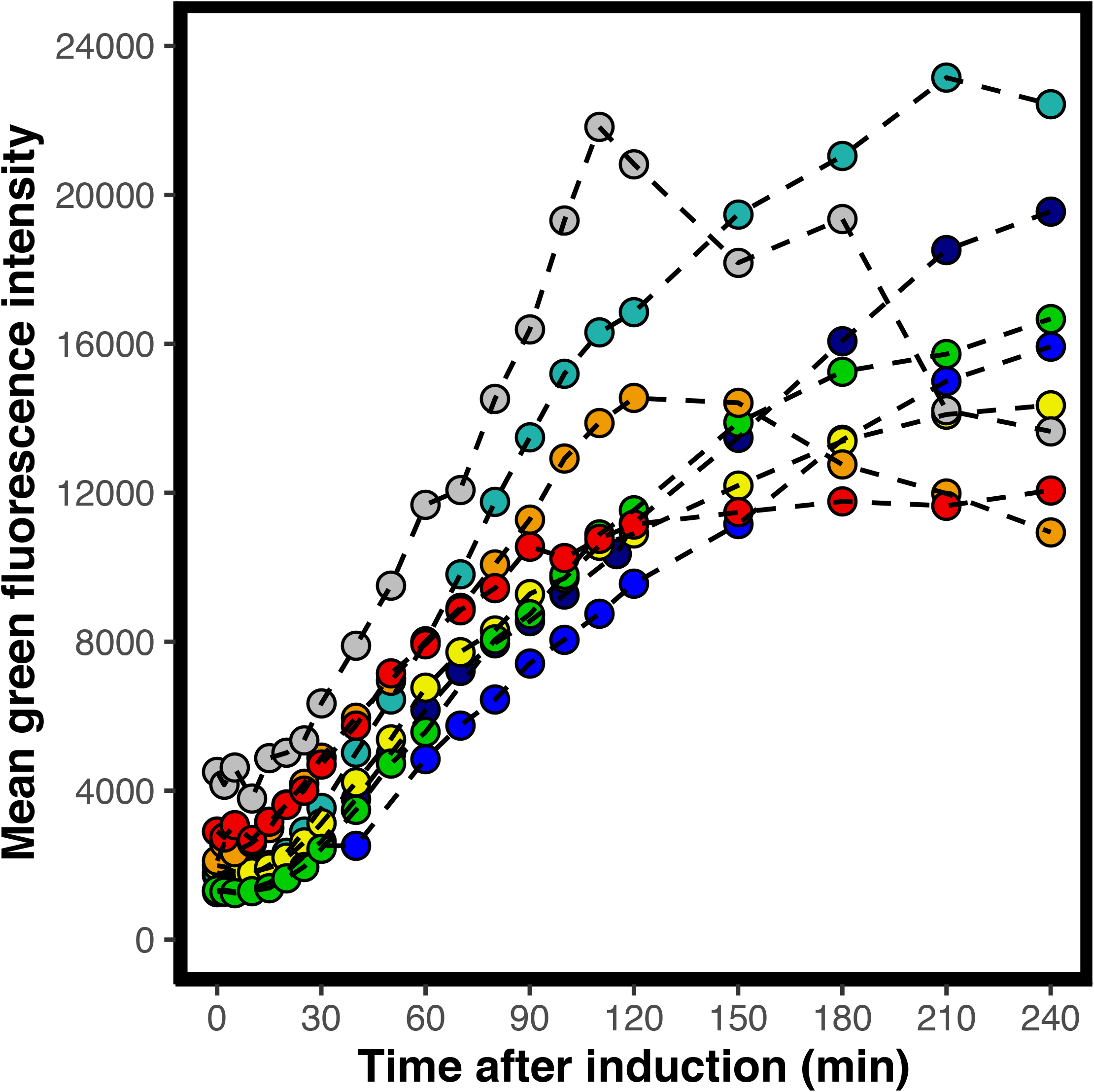
Transcription from the DnaK promoter monitored by green fluorescence. Mean green fluorescence intensity of cells expressing variants: L57G (dark blue), L57C (blue), L57P (cyan), M40A (green), Q33S (yellow), V36I (orange), Q33Y (red) and wt (grey). One of two replicates are shown for each variant.

**S 5.**
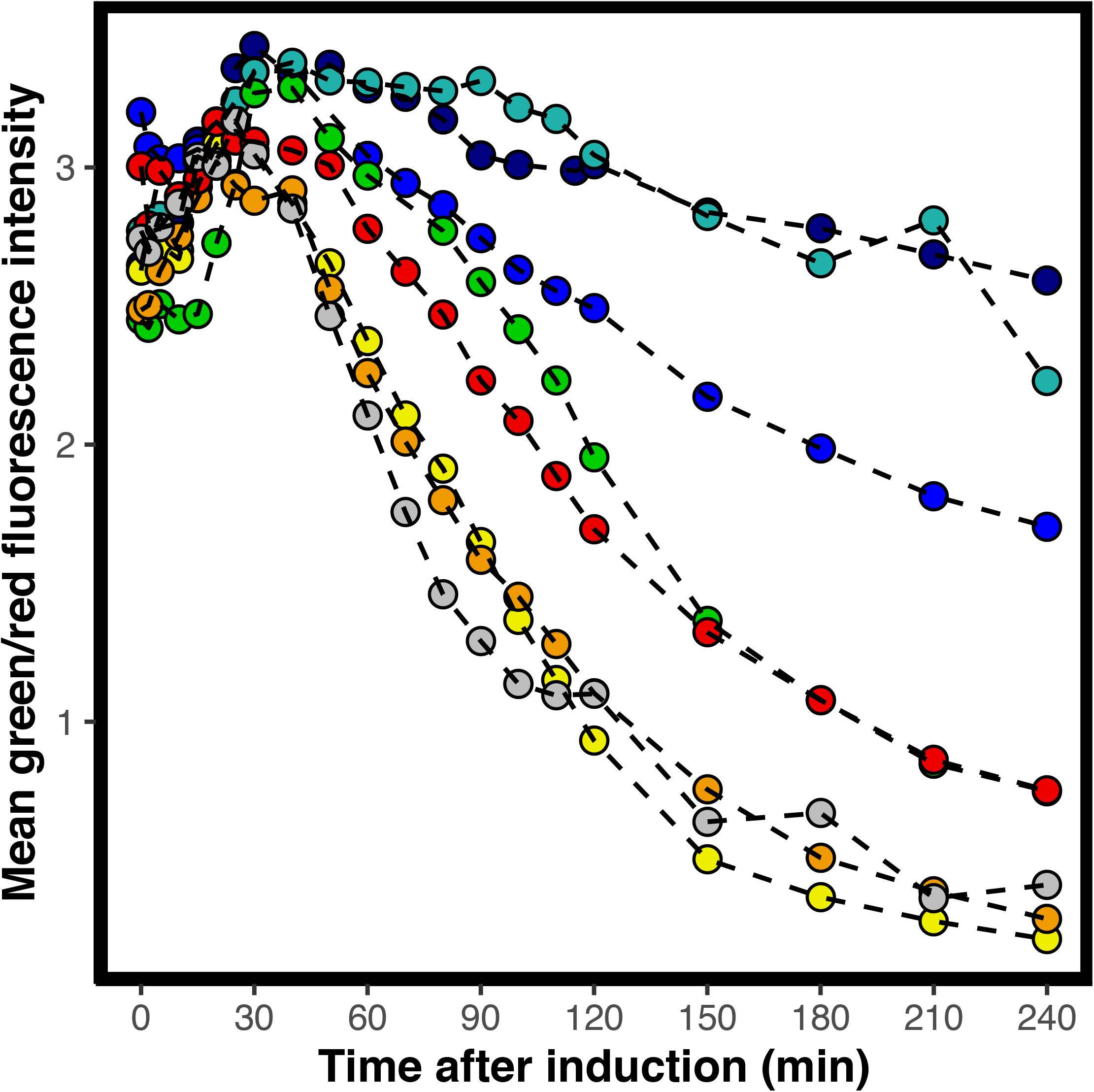
Accumulated transcription from the DnaK promoter normalised to protein abundance. Mean green fluorescence intensity divided by red fluorescence intensity of cells expressing variants: L57G (dark blue), L57C (blue), L57P (cyan), M40A (green), Q33S (yellow), V36I (orange), Q33Y (red) and wt (grey). One of two replicates are shown for each variant.

**S 6.**
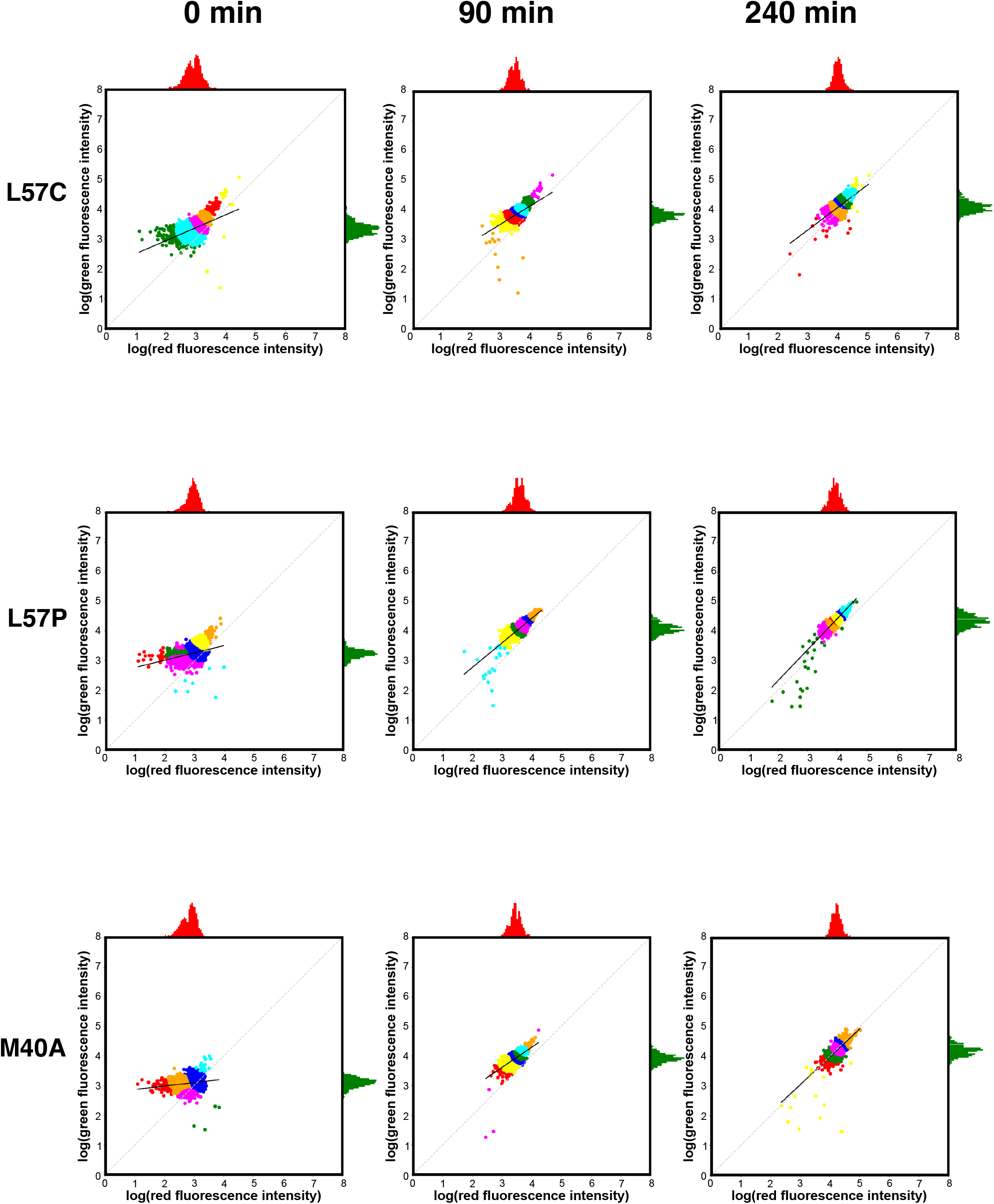
Red and green fluorescence of single cells expressing L57C, L57P and M40A. The red and green fluorescence for each individual cell in a population expressing L57C (top), L57P (middle) and M40A (bottom) at the time of induction (left) and 240 min after induction with IPTG (right). Colouring is based on a Bayesian Gaussian mixture model for cluster assignment. The x-axis histogram (red) shows the distribution of red fluorescence, and the y-axis histogram (green) shows the distribution of green fluorescence within the population.

**S 7.**
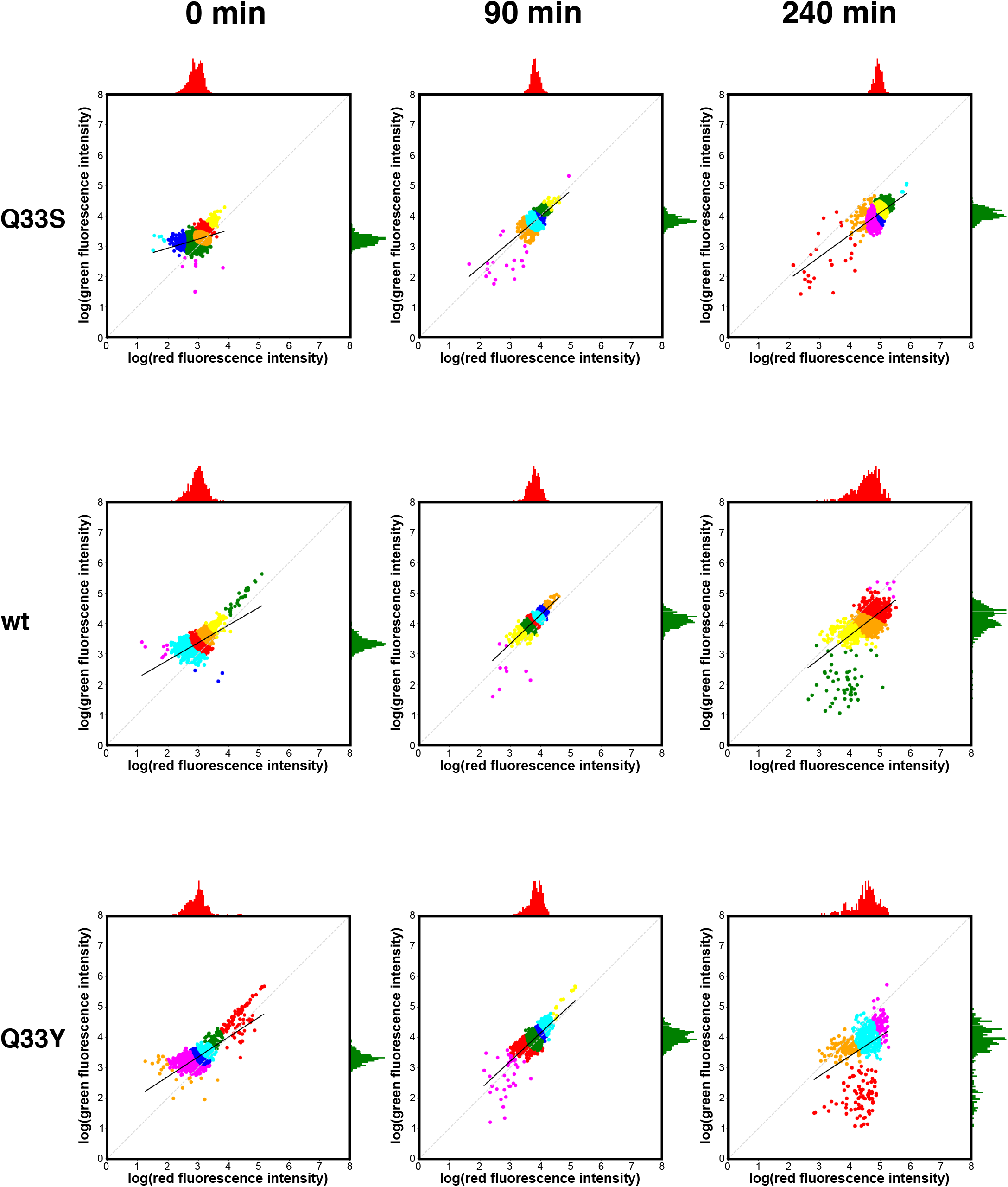
Red and green fluorescence of single cells expressing Q33S, wt and Q33Y. The red and green fluorescence for each individual cell in a population expressing Q33S (top), wt (middle) and Q33Y (bottom) at the time of induction (left) and 240 min after induction with IPTG (right). Colouring is based on a Bayesian Gaussian mixture model for cluster assignment. The x-axis histogram (red) shows the distribution of red fluorescence, and the y-axis histogram (green) shows the distribution of green fluorescence within the population.

**S 8.**
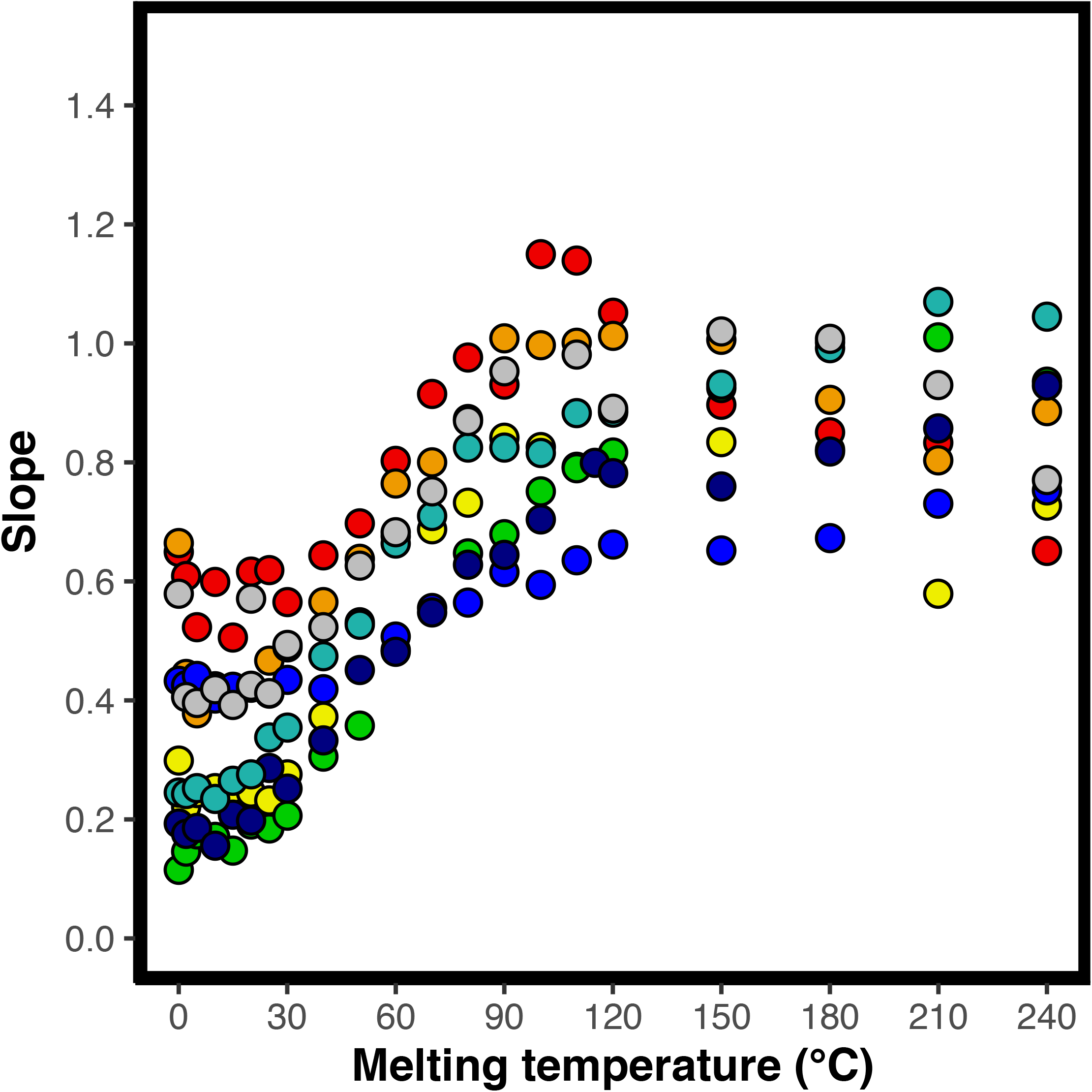
Slope of fluorescent population over time. For a population of cells expressing the variants: L57G (dark blue), L57C (blue), L57P (cyan), M40A (green), Q33S (yellow), V36I (orange), Q33Y (red) and wt (grey) the slope of log transformed red fluorescence vs log transformed green fluorescence is calculated. One of two replicates are shown for each variant.

**S 9.**
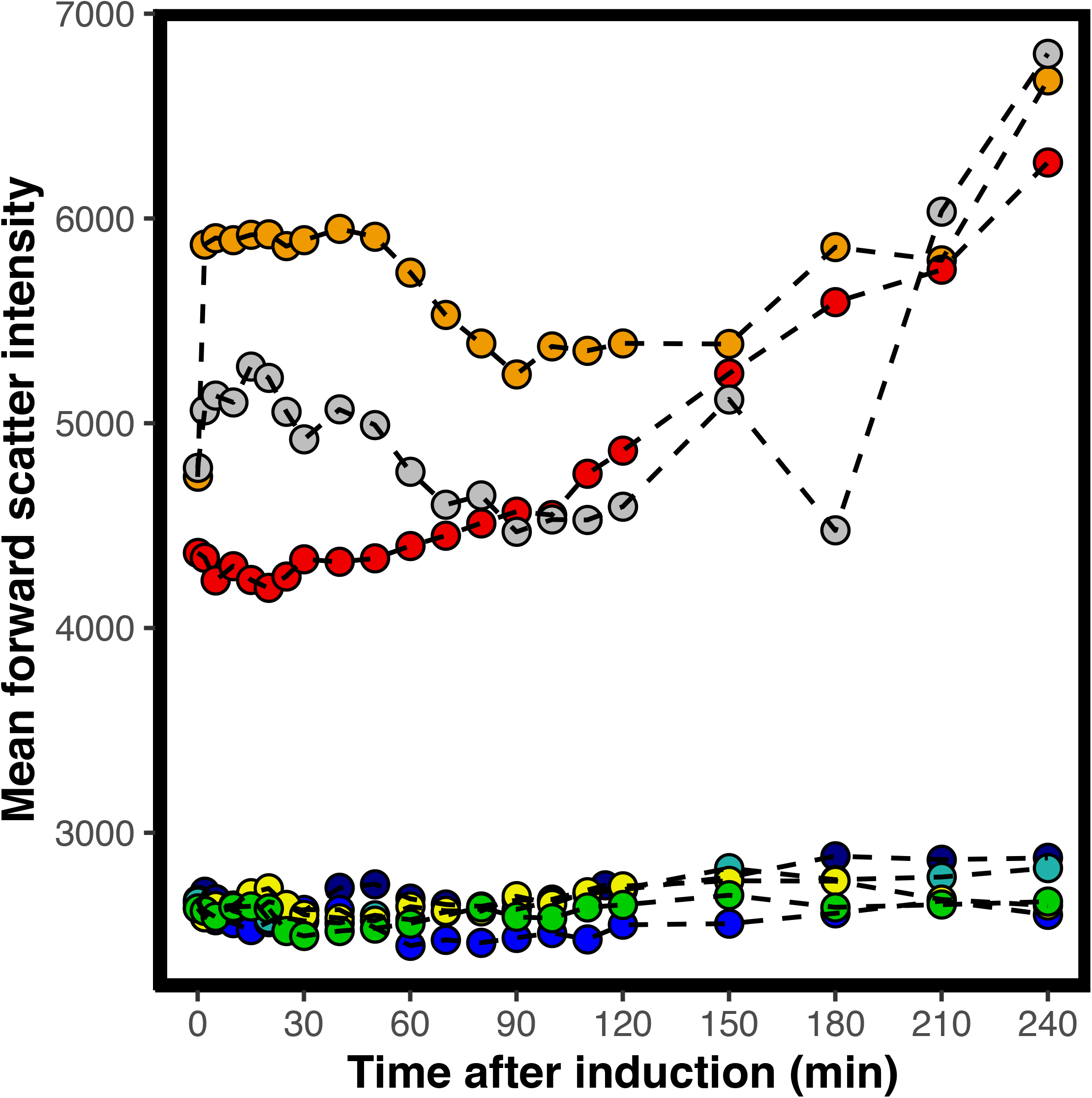
Forward scatter of whole populations expressing N102LT variants. Forward scatter as a function of time for variants: L57G (dark blue), L57C (blue), L57P (cyan), M40A (green), Q33S (yellow), V36I (orange), Q33Y (red) and wt (grey) One of two replicates are shown for each variant.

**S 10.**
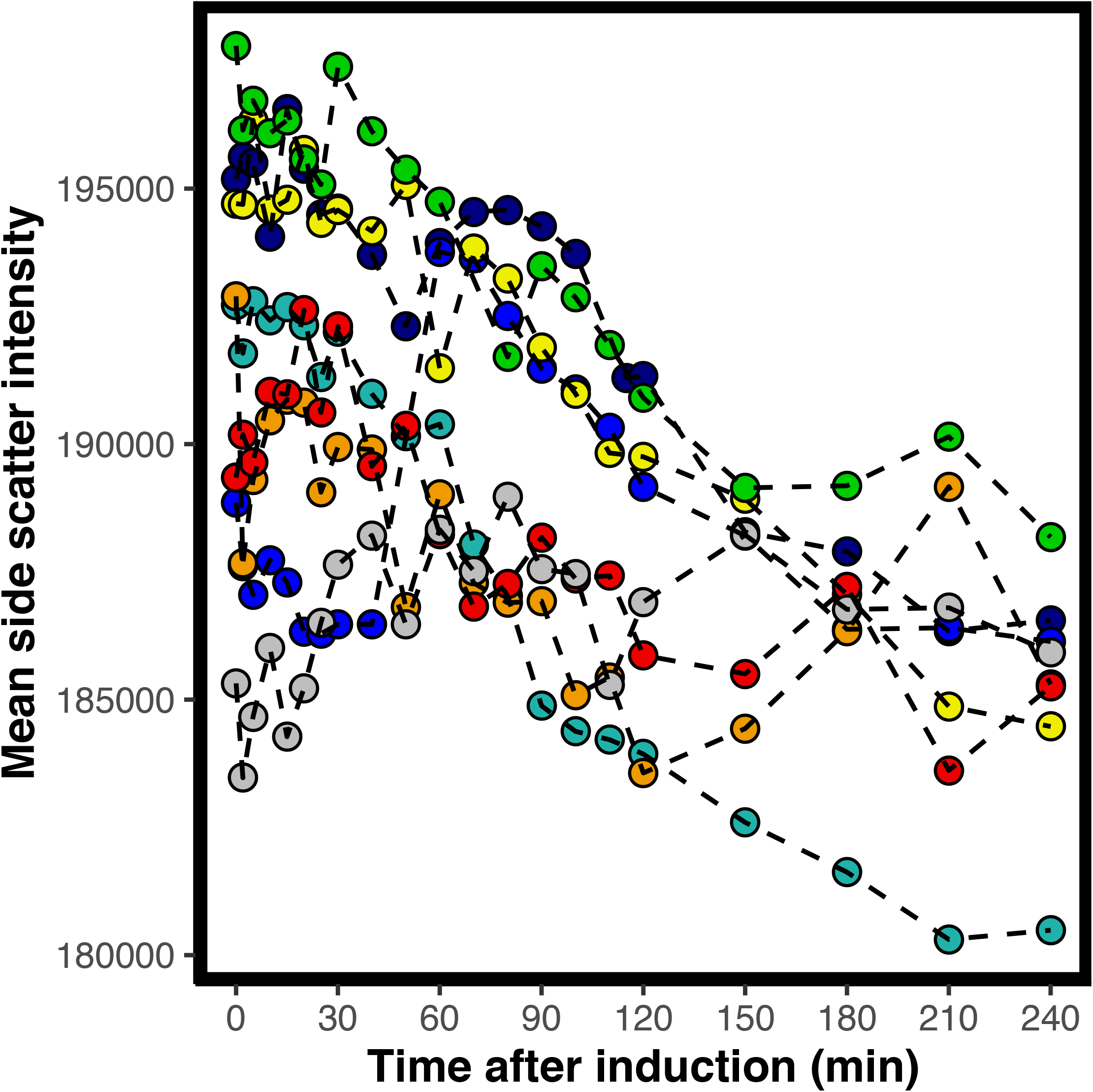
Side scatter of whole populations expressing N102Lt variants. Side scatter as a function of time for variants: L57G (dark blue), L57C (blue), L57P (cyan), M40A (green), Q33S (yellow), V36I (orange), Q33Y (red) and wt (grey). One of two replicates are shown for each variant.

